# Hydrophobic mismatch induces lipid sorting based on tail unsaturation

**DOI:** 10.64898/2026.06.23.734047

**Authors:** Niek van Hilten, Michael Grabe

**Affiliations:** Department of Pharmaceutical Chemistry, University of California, San Francisco, San Francisco, California, 94158, United States of America; Cardiovascular Research Institute, University of California, San Francisco, San Francisco, California, 94158, United States of America

## Abstract

Biological membranes contain a diverse set of membrane proteins surrounded by many different lipids, and the lateral organization and function of these molecules are closely intertwined. Here, we use coarse-grained molecular dynamics (MD) simulations to explore how hydrophobic mismatch between the length of transmembrane (TM) proteins and the thickness of the surrounding lipid membrane impacts the spatial distribution of the lipids. We constructed idealized cylindrically symmetric proteins, inspired by the “Mattress Model” developed in the 1980’s, and simulated these model proteins in different lipid compositions. We found that unsaturated lipids were attracted to *short* TM proteins that thinned the membrane, while fully saturated lipids were attracted to *long* TM proteins that induced membrane extension. A simple mechanical description of the membrane deformation energy coupled to a lipid mixing model accurately predicted the enrichment/depletion, which was up to 33% in some cases. Our simulations also highlight that lipid sorting behavior is sensitive to protein tilt and protein surface roughness. By teasing out the fundamental physical principles in these simple models, our results provide a foundational understanding of how proteins and lipids form complex and transient assemblies, which we believe will be important for interpreting lipid-protein interactions for a host of membrane proteins that regulate cellular membranes and cell function.

**SIGNIFICANCE:** Transmembrane (TM) proteins in the eukaryotic plasma membrane are solvated by hundreds of lipid species with different head group chemistries and tail configurations. This work explores how the number of double C-C bonds in the lipid acyl chains (unsaturations) affects the enrichment and depletion of lipid species around TM proteins that exhibit hydrophobic mismatch: i.e., they are either too long or too short for the membrane. We use molecular simulation and elastic modeling to show that short TM proteins attract unsaturated lipids, while long TM proteins attract saturated ones. This finding unlocks a currently understudied aspect of protein-lipid interactions that has important implications for our fundamental understanding of membrane proteins.

## INTRODUCTION

The lipid and protein contents of biological membranes are characterized by tremendous complexity. The bilayers that encapsulate eukaryotic organelles have distinct lipid compositions and thicknesses (1), features that are related to the proteins embedded in them (2, 3). Over the last several decades, researchers have extensively studied how discrepancies between the size of transmembrane (TM) proteins and the surrounding lipids (a phenomenon called “hydrophobic mismatch”) give rise to biophysical driving forces that govern membrane-mediated protein aggregation (4–8), proteins partitioning into specific lipid environments (9, 10), and the association of specific lipid species to the protein-membrane interface to form a proteolipid fingerprint (1, 11–13).

One of the most influential physical descriptions of hydrophobic mismatch is the “mattress model” published by Mouritsen and Bloom in 1984 (14). They approximated transmembrane proteins and the surrounding bilayer environment by a system of parallel springs, each with their respective characteristic equilibrium length and spring constant. Consequently, the membrane shape around TM proteins that are too long or too short for their lipid environment follows from a tug of war between the elastic energies of extending/compressing surrounding lipid molecules and the exposure of the apolar protein surface to the water phase for oversized proteins or membrane burial of polar amino acids for short proteins. Follow-up studies showed that the mattress model can be used to predict how lipids with different acyl chain lengths sort to or from protein inclusions to minimize hydrophobic mismatch (15, 16).

Here, we constructed simplified protein-lipid model systems to address the question of whether lipid tail (un)saturation alone is sufficient to govern lipid sorting behavior with respect to hydrophobically mismatched TM proteins. Inspired by the mattress model, we performed coarse-grained molecular dynamics (CGMD) simulations of idealized cylindrical “pseudoproteins” with different lengths in two-component lipid mixtures. Based on the well-established phenomenon that increasing the number of double bonds in lipid tails decreases bilayer thickness, increases lipid disorder, and softens the membrane (17, 18), we hypothesized that unsaturated lipids would accumulate around short TM proteins to help relieve the elastic energy of the membrane. Following the same argument, we predicted that saturated lipids would enrich near long TMs. Strikingly, our CGMD simulations indeed show the expected sorting trends.

Finally, we formulated an elastic membrane model that mirrors our CGMD setups and applied a Markov-chain Monte Carlo algorithm to find the optimal lipid distribution that minimizes the system’s free energy. We parametrized this model with experimentally determined physical lipid parameters for lipids with 0, 1, 2, or 4 unsaturations from the literature and found that predicted lipid distributions accurately match the lipid sorting trends we observed in our CGMD simulations.

Taken together, our findings underline the importance of lipid tail configuration, and how something as subtle as a double C-C bond may affect the local membrane composition and functionality of transmembrane proteins with different lengths.

## METHODS

### Cylindrical pseudoproteins

Martini models of cylindrical “pseudoproteins” were constructed by fitting a triangular grid of beads with grid spacing *b* to a circle with a 25 Å radius (Fig. S1a,b), approximately matching the typical size of a transmembrane protein. These circular arrangements were then stacked with a uniform *z* = 5 Å spacing (Fig. S1c) to form cylinders with total heights of 20 Å (C20, 5 layers), 30 Å (C30, 7 layers), and 40 Å (C40, 9 layers). All beads in the top and bottom layers were assigned a hydrophilic P4 bead type. The middle layers all consisted of hydrophobic C3 beads. We deliberately opted for C3 beads because they have the same Lennard-Jones parameters for their non-bonded interactions with saturated lipid tail beads (type C1) and unsaturated lipid tail beads (type C4h) within the Martini 3.0.0 force field (19). In doing so, we aimed to eliminate the role of direct protein-lipid attraction/repulsion in our simulations, such that any lipid sorting can be attributed mostly to membrane elastic effects. The bond length 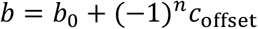 between the beads within a single layer (labeled *n*) is alternated with an offset factor *c*_offset_to control the roughness of the pseudoprotein surface (Fig. S1d). In all simulations, *b*_0_ = 4.7 Å and *c*_offset_ = 0.1 Å, unless noted otherwise. All beads within 9 Å of each other were interconnected by an elastic network with a force constant *k*= 500 kJ mol^−1^nm^−2^.

### CGMD simulations

Pseudoproteins were embedded in membranes using the *insane* python script (20) (Fig. 1a, Table S1 for all setups). Two-component membranes all contained 70 mol% of 1,2-dioleoyl-sn-glycero-3-phosphitadylcholine (DOPC, 18:1-18:1) as a “host” lipid and 30 mol% of variable “guest” lipids that were placed randomly in the initial system. As guest lipids, we chose 1,2-distearoyl-sn-glycero-3-phosphitadylcholine (DSPC, 18:0-18:0), 1-palmitoyl-2-oleoyl-sn-glycero-3-phosphitdaylcholine (POPC, 16:0-18:1), 1-palmitoyl-2-linoleoyl-sn-glycero-3-phosphitdaylcholine (PLiPC, 16:0-18:2), and 1,2-dilinoleoyl-sn-glycero-3-phosphitadylcholine (DLiPC, 18:2-18:2). Throughout, we use X:Y notation where X is the number of carbons and Y is the number of double bonds in the acyl chain. All simulations were performed using the Gromacs MD engine, version 2023.3, (21) with the Martini force field, version 3.0.0 (19), in which DSPC, PLiPC, and DLiPC are parametrized and abbreviated as DPPC, PIPC, and DIPC, respectively. After energy minimization and a brief NPT equilibration (1 ns, using the c-rescale barostat (22)), we performed 10 μs simulations for each lipid mixture. We used a 20 fs time step and a 0.5 ns^−1^ save rate. Unless noted otherwise, we applied a tilt angle restraint potential (with force constant *k*= 10,000 *kJ mol*^−1^*rad*^−2^) to maintain a 0° angle between the z-axis and the vector between the centers-of-mass of the two hydrophilic layers of the pseudoproteins. We kept our systems at a constant temperature of 323 K (τ_*T*_ = 1 *ps*) using the v-rescale thermostat (23) and applied semi-isotropic Parrinello-Rahman pressure coupling (24) to maintain a 1.0 bar pressure in all directions (τ_*P*_ = 12 *ps*, compressibility = 3 × 10^−4^ bar^−1^). A 1.1 nm cut-off distance was used for reaction-field electrostatics and Van der Waals potentials (25), and the Verlet scheme was applied for updating the neighbor list every 20 steps with a 1.35 nm cut-off (26).

**Figure 1.**
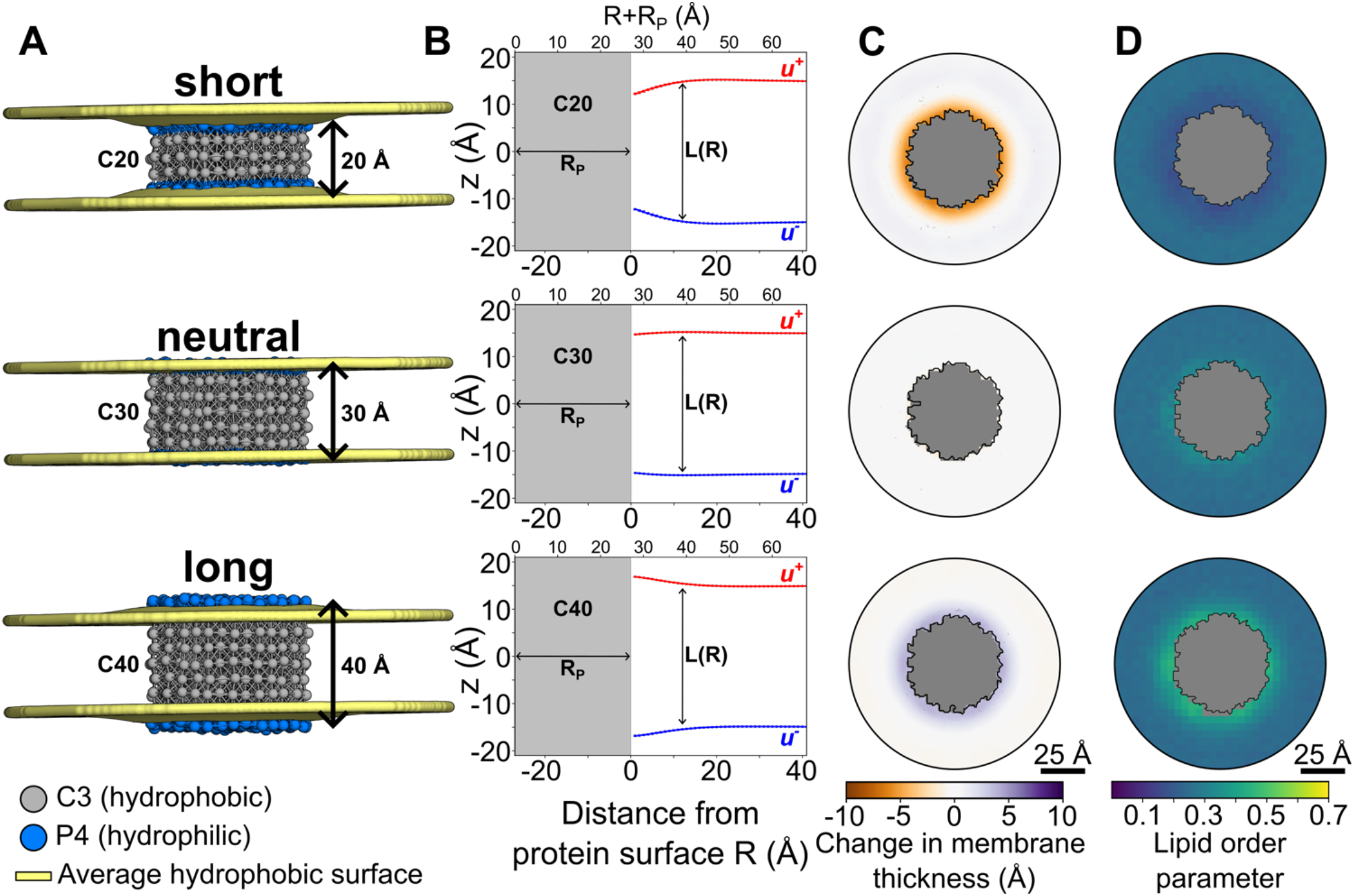
Pseudoproteins induce membrane deformations and change lipid order in pure DOPC (18:1,18:1) bilayers. **(a)** Side views of the C20, C30, and C40 pseudoproteins simulated in a pure DOPC bilayer. Yellow surfaces represent the ensemble-averaged glycerol positions (GL1 and GL2 beads). **(b)** Average leaffet shapes from CGMD simulation, fit with C^th^ degree polynomials representing u^+^ and u^−^ (Table S2). The vertical distance between the lines yields the hydrophobic thickness L(R) = u^+^ − u^−^. **(c)** Top view membrane thickness maps based on the yellow surfaces in panel a. Gray represents the pseudoprotein. **(d)** Lipid order parameter maps around pseudoproteins in pure DOPC. Gray represents the pseudoprotein.

### CGMD simulation analysis

Prior to all analyses, we centered and aligned all simulation trajectories with respect to the pseudoprotein coordinates in the initial frame, using the Gromacs tool *gmx trjconv*. In all plots, pseudoprotein shapes were constructed by drawing a concave hull (27) around the aggregated xy-coordinates (plus Van der Waals radii) from 50 evenly spaced snapshots. We excluded the first 1 μs from all analyses for equilibration, and applied three analysis protocols:

#### 1. Membrane shape

The average membrane shape was calculated from the glycerol positions (GL1 and GL2 beads) of each lipid, following a protocol described previously (28, 29). In brief, we applied linear interpolation to a rectilinear grid with a uniform 1 Å spacing and averaged over all simulation frames (1-10 μs). Scarcely occupied grid points (<2% occupancy) were discarded. The local membrane thickness was calculated as the height difference between the two resulting ensemble-averaged surfaces (upper and lower). Data were then averaged radially around the protein and fit with a 6^th^ degree polynomial (Table S2) to yield upper and lower leaflet shapes *u*^+^ and *u*^−^, respectively.

#### 2. Lipid order parameter

For each frame, lipids were assigned to the upper or lower leaflet based on the z-coordinate of the phosphate (PO4). We constructed a rectilinear grid and assigned each lipid in each frame to 4×4 Å^2^ bins according to the xy-coordinates of the phosphate (PO4) bead. For each tail bond in each lipid in each frame, we calculated the angle *θ* between the bond vector and the local leaflet surface normal based on the ensemble-averaged surface calculated in the previous analysis step. Per bin, these angles were used to compute the lipid order parameter:

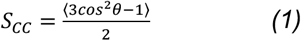

Data were averaged over all frames and both leaflets to yield 2D lipid order maps.

#### 3. Lipid molar fractions

In the two-component lipid mixtures, we collapsed the two leaflets onto the xy-plane and calculated the time-cumulated occurrence of phosphate (PO4) beads in 2×2 Å^2^ bins for the host (*n*_*h*_) and the guest lipid (*n*_g_) using MDAnalysis (30). Then, we calculated the host and guest lipid molar fractions as 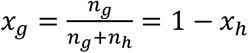 in each bin and plotted them in 2D maps.

Radial averages and standard deviations were calculated by constructing radial isosurface bins with a 1 Å spacing around the pseudoprotein shape using the *shapely* python module (31) (Fig. S2).

### Elastic membrane model with lipid mixing

To better understand the energetics behind our CGMD simulation results, we defined a continuum elastic model of cylindrical pseudoproteins C20, C30, and C40 with radius *R*_*P*_ embedded in two-component lipid mixtures with 30 mol% of variable guest lipid (labeled *g*) and 70 mol% of DOPC host lipid (labeled *h*). In all cases, we use the ensemble-averaged leaflet shapes *u*^+^ and *u*^−^ from our CGMD simulations of pure DOPC around the respective pseudoproteins (C20, C30, C40, see Fig. 1b). Our model includes a bilayer compression term *H*_comp_, a leaflet bending term *H*_bend_, and a mixing entropy term *S*_mix_ to define the system’s free energy as:

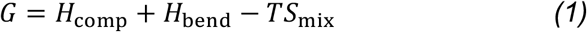

#### 1. Compression

We consider the bilayer compression *H*_comp_ as proposed by Huang (32) and the original mattress model (14). That is, we assume that each lipid type has a preferred equilibrium length *L*_0,g_ and *L*_0,*h*_ and that the energy grows quadratically in the strain as the bilayer deviates from this value, which is calculated from the length *L*(*R*) = *u*^+^ − *u*^−^, i.e., the instantaneous bilayer thickness at distance *R* from the protein surface in a deformed membrane with upper and lower leaflet shapes *u*^±^. Thus, per unit area, we have:

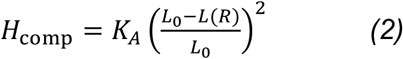

Next, given the characteristic area compressibility moduli *K*_*A*,g_ and *K*_*A,h*_ for the guest and host lipid, respectively, we assume that the relative contribution to the energy from each lipid species is linearly proportional to its mole fraction, *x*_g_ or *x*_*h*_. For each discrete radial bin *j* with width *d* and area *A*_j_ = 2*πd*(*R*_j_ + *R*_*P*_) at distance *R*_j_ from the protein surface, we write:

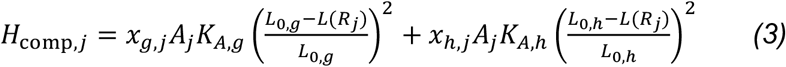

where for example *x*_g,j_ = *x*_*lipidtype,bin*_.

#### 2. Bending

The leaflet bending energy 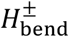 is calculated from the MD-derived leaflet shapes *u*^+^ and *u*^−^ as done by others (32–34). In cylindrical coordinates, we define the leaflet shapes *u*^±^ to be a function of *r* = *R*_j_ + *R*_*P*_, which is the radial distance from the origin to the center of a bin *j*. Since the system is cylindrically symmetric, we can write:

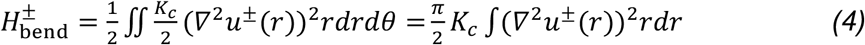

where *K*_*c*_ is the bilayer bending modulus that it is divided by 2 to account for a single leaflet. Considering distinct bending moduli for each lipid species we have *K*_*c*,g_ and *K*_*c,h*_, fractions *x*_g_ or *x*_*h*_, and bin width *d* we write for each bin *j*:

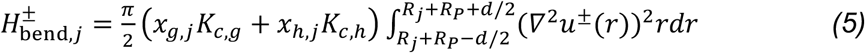

Finally, we sum the contributions of the upper and lower leaflet together to get 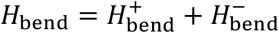 hat goes into Eq. 1.

#### 3. Mixing entropy

The third and final term in the free energy is the mixing entropy. For our guest and host lipids in discrete bins *j*, this follows the text-book definition (35):

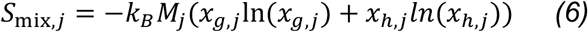

where *k*_*B*_ is the Boltzmann constant, and *M*_j_ is the number of molecules which we get from the bin area *A*_j_ as *M*_j_ = 2*A*_j_⁄*A*_lip_. Here, the factor two comes from the two membrane leaflets and *A*_j_ = 2*πd*(*R*_j_ + *R*_*P*_) is the membrane area of the bin. We assume that the area per lipid *A*_lip_ is a linear combination of the characteristic areas per lipid for the guest and host lipid at constant bulk concentrations *C*_g_ = 0.3 and *C*_*h*_ = 0.7: *A*_lip_ = *C*_g_*A*_lip,g_ + *C*_*h*_*A*_lip,*h*_.

#### 4. Markov-chain Monte Carlo optimization

Next, we set out to find the radial lipid distributions *x*_g_ and *x*_*h*_ = 1 − *x*_g_ that minimize the free energy difference Δ*G* with respect to a uniform 3:7 distribution. We divided our system into *N* radial bins labelled *j* ∈ {1, …, *N*}, with a distance to the protein surface *R*_j_ and bin width *d* that we set to 1 Å (Fig. 2). We start at step *i* = 0 with a homogeneously mixed system where the lipid ratio is 3:7 everywhere, i.e., *x*_g_ = *C*_g_ = 0.3 and *x*_*h*_ = *C*_*h*_ = 0.7. To minimize the system’s free energy, we applied a Markov-chain Monte Carlo (MCMC) optimization scheme that iteratively makes small, random, changes in the guest lipid fractions *x*_g,*i*,j_ in all bins *j* simultaneously, where for clarity *x*_g,*i*,j_ = *x*_*lipidtype,timestep,bin*_ and the time index has been inserted between lipid type and radial bin number. The mole fraction of host lipid in each bin is adjusted accordingly by *x*_*h,i*,j_ = 1 − *x*_g,*i*,j_. Along the same lines, we write the *number of lipids* in each bin as *n*_g,*i*,j_ = *n*_*lipidtype,timestep,bin*_. The total lipid number in each bin (*n*_g,*i*,j_ + *n*_*h,i*,j_) is kept constant at a value *M*_j_ that is determined by the area of the bin and the area per lipid, following *M*_j_ = 2*A*_j_⁄*A*_lip_, as described in the previous section.

**Figure 2.**
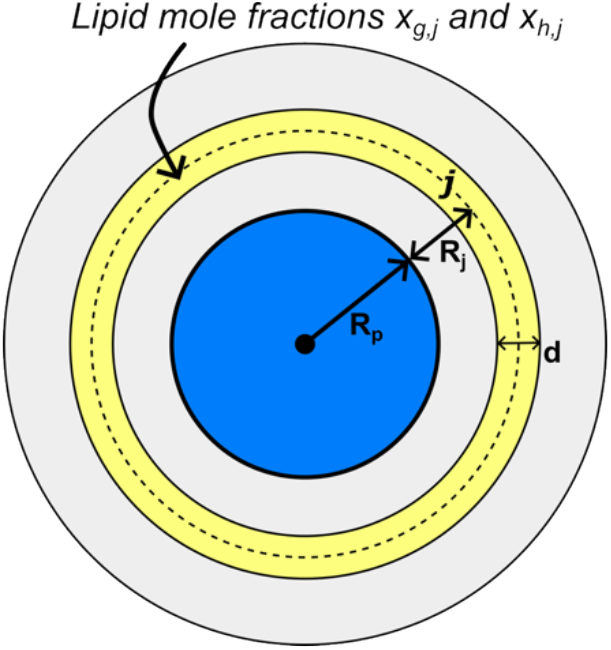
MCMC optimization setup. The blue circle depicts a top view of a pseudoprotein with radius R_p_. The yellow radial bin j has width d = 1 Å (not drawn to scale) and its area is given by A_j_ = 2πd(R_j_ + R_P_). The total number of lipids (guest plus host) in each bin is 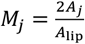

At each update step *i*, we converted the list of guest lipid *fractions* per bin from the previous step *x*_g,*i*–1,j_ to the guest lipid *numbers n*_g,*i*–1,j_ by multiplying each mole fraction by the total number of lipids in the respective bin *n*_g,*i*–1,j_ = *x*_g,*i*–1,j_*M*_j_. For bins *j* ∈ {1, …, *N* − 1}, we perturb the number of lipids by adding a random number sampled from a uniform distribution between −0.01 and 0.01: *α*_*rand*,j_ = *U*{−0.01, 0.01}. For the last bin (*j* = *N*), we assign a final value 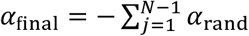 to yield the “balanced” update list (*α*_*rand*,1_, …, *α*_*rand,N*–1_, *α*_*final*_) that sums to zero 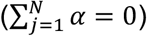 ensuring that the total number of lipids in the system is always constant. Furthermore, we rejected and regenerated update lists if | *α*_*final*_ | > 0.01 or if *x*_g,*i*_ > 1 so that the total guest lipid fraction does not exceed 1 in any bin. The new lipid numbers are converted back to mole fractions by dividing by the total lipid number density everywhere *x*_g,*i*,j_ = *n*_g,*i*,j_/*M*_j_. Then, we calculate the total free energy difference between this updated lipid distribution and the initial uniform state (*x*_g,*i*=0_ = *C*_g_ = 0.3 everywhere) using Eq. 1 as follows:

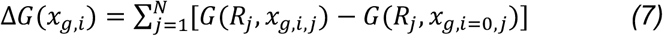

Monte Carlo moves *x*_g,*i*–1_ → *x*_g,*i*_ are accepted or rejected using a Metropolis criterion where the acceptance probability is given by:

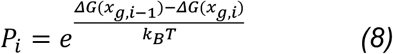

We included a simplified numerical example of the entire updating scheme in Table S3.

#### 5. Model parametrization and settings

Ten independent MCMC optimization runs with 5,000 iterations were performed for each guest lipid type DSPC, POPC, PLiPC, and DLiPC. We parametrized the elastic membrane model with each lipid’s equilibrium length *L*_0_, area compressibility modulus *K*_*A*_, bilayer bending modulus *K*_*c*_, and area per lipid *A*_lip_. These parameters were derived from experimental values and atomistic MD simulation data from the literature (Table 1).

**Table 1.**
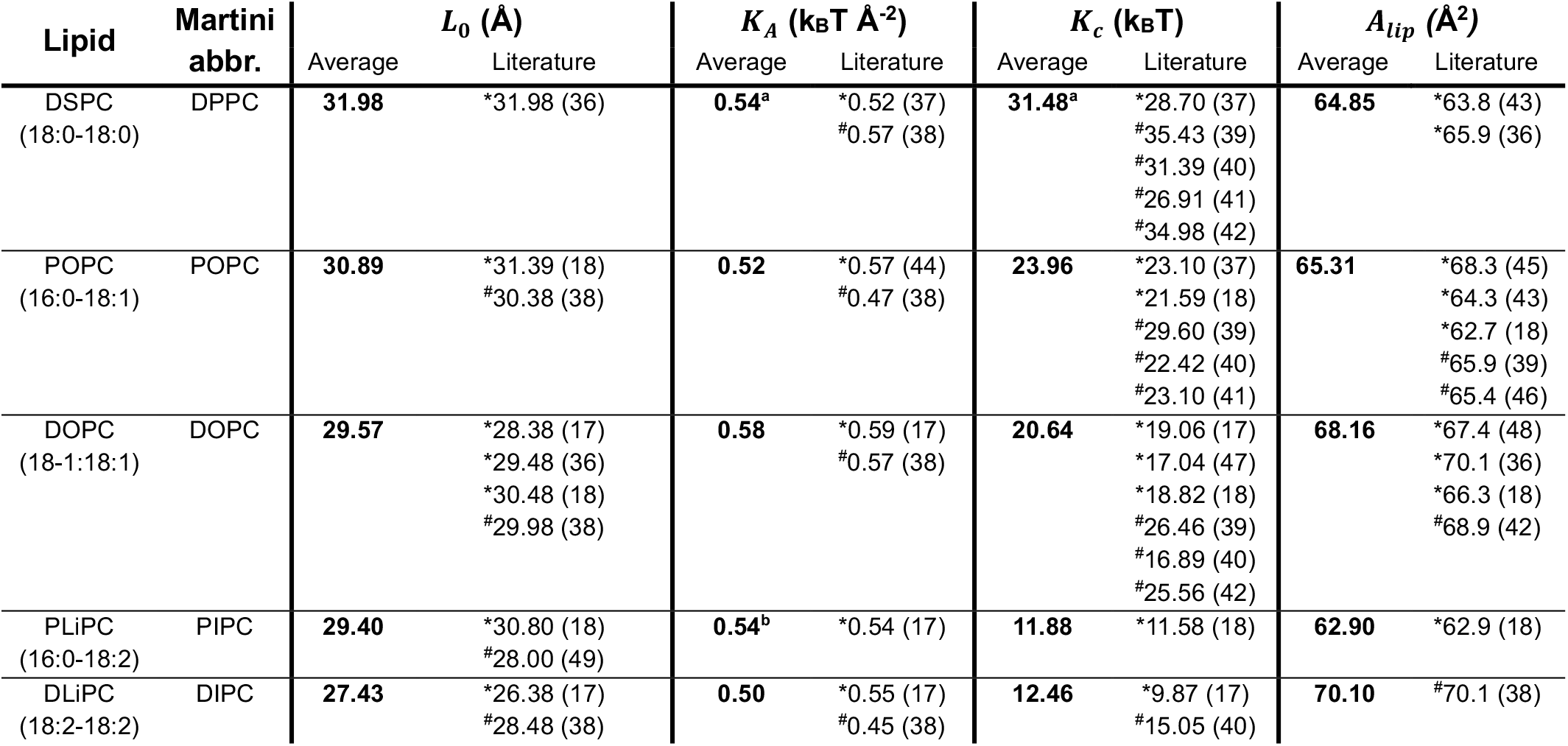
Physical lipid parameters. Final values are printed in bold and represent the averages of literature values. We obtained equilibrium lipid lengths L_0_ = h_pp_ − h_0_ by taking membrane thicknesses h_pp_ from the literature and subtracting the thickness of the incompressible headgroup region from CGMD simulations h_0_=8.52 Å. If applicable, moduli were converted to units of k_B_T with T = 323 K to match the CGMD simulation settings. *Experiment; ^#^All-atom simulation; ^a^values for DPPC (16:0-16:0), in lack of DSPC data; ^b^values for SLiPC (18:0-18:2), in lack of PLiPC data.

## RESULTS

### Idealized protein inclusions deform and disorder membranes

To assess how protein-membrane hydrophobic mismatch governs (un)saturation-based lipid sorting, we constructed three cylindrical “pseudoproteins” with hydrophobic thicknesses of 20, 30, and 40 Å (named C20, C30, and C40; see Fig. 1a). For the Martini 3.0.0 force field, we measured an average hydrophobic thickness of 29 Å for a protein-free DOPC bilayer (Table S4), which means that C20 is too short for the membrane, C30 is neutral, and C40 is too long for the membrane. We embedded each pseudoprotein in a pure DOPC bilayer and performed 10 μs of CGMD simulation to determine the ensemble-averaged membrane shape around the protein (Fig. 1a-c). For the short C20 protein, we observed 5.4 Å of bilayer thinning (18%) at the protein-membrane interface at *R* = 1 Å, compared to the hydrophobic thickness far away from the protein at *R* = 40 Å. In contrast, the long C40 protein locally increased the hydrophobic thickness by 3.9 Å (13%). C30, designed to match the bilayer equilibrium thickness, indeed resulted in minimal deformations, thinning the membrane by only 0.5 Å (2%) at *R* = 1 Å. We also measured the average lipid order parameter around the pseudoproteins and found that C20 *decreases* and C40 *increases* the order of the surrounding lipid tails in pure DOPC (Fig. 1d). This is a direct result of compressing (C20) and extending (C40) the lipid molecules to match the hydrophobic thickness of the pseudoproteins.

### Protein-induced membrane deformations govern lipid sorting based on tail saturation

Next, we repeated the simulations in two-component lipid mixtures. With a constant background of 70 mol% DOPC (18:1-18:1) as the host lipid (*C*_*h*_ = 0.7) we varied the degree of unsaturation in the remaining 30 mol% of guest lipid (*C*_g_ = 0.3) and considered DSPC (18:0-18:0), POPC (16:0-18:1), PLiPC (16:0-18:2), and DLiPC (18:2-18:2). Membrane thickness profiles of the mixed membranes all showed the same shapes as pure DOPC (Fig. S3), with subtle (~2 Å) uniform thickening for the more saturated DSPC/DOPC and POPC/DOPC mixtures and similar thinning for more unsaturated PLiPC/DOPC and DLiPC/DOPC mixtures. For each simulation, we aligned and centered the trajectory on the starting coordinates of the pseudoprotein and calculated ensemble-averaged guest lipid concentration maps (Fig. 3a). We used 1 Å isosurface bins at increasing distances R from the surface of the pseudoprotein and applied block averaging (1-4, 4-7, 7-10 μs) to quantify the average guest lipid mole fraction at discrete intervals (Fig. S2). This analysis revealed that the short C20 protein attracted unsaturated lipid species and repelled fully saturated lipids (Fig. 3b). Conversely, the vicinity of the long C40 was enriched in saturated lipids and depleted in unsaturated lipids (Fig. 3b). The neutral C30 did not cause significant enrichment/depletion in any of the four lipid mixtures we tested.

**Figure 3.**
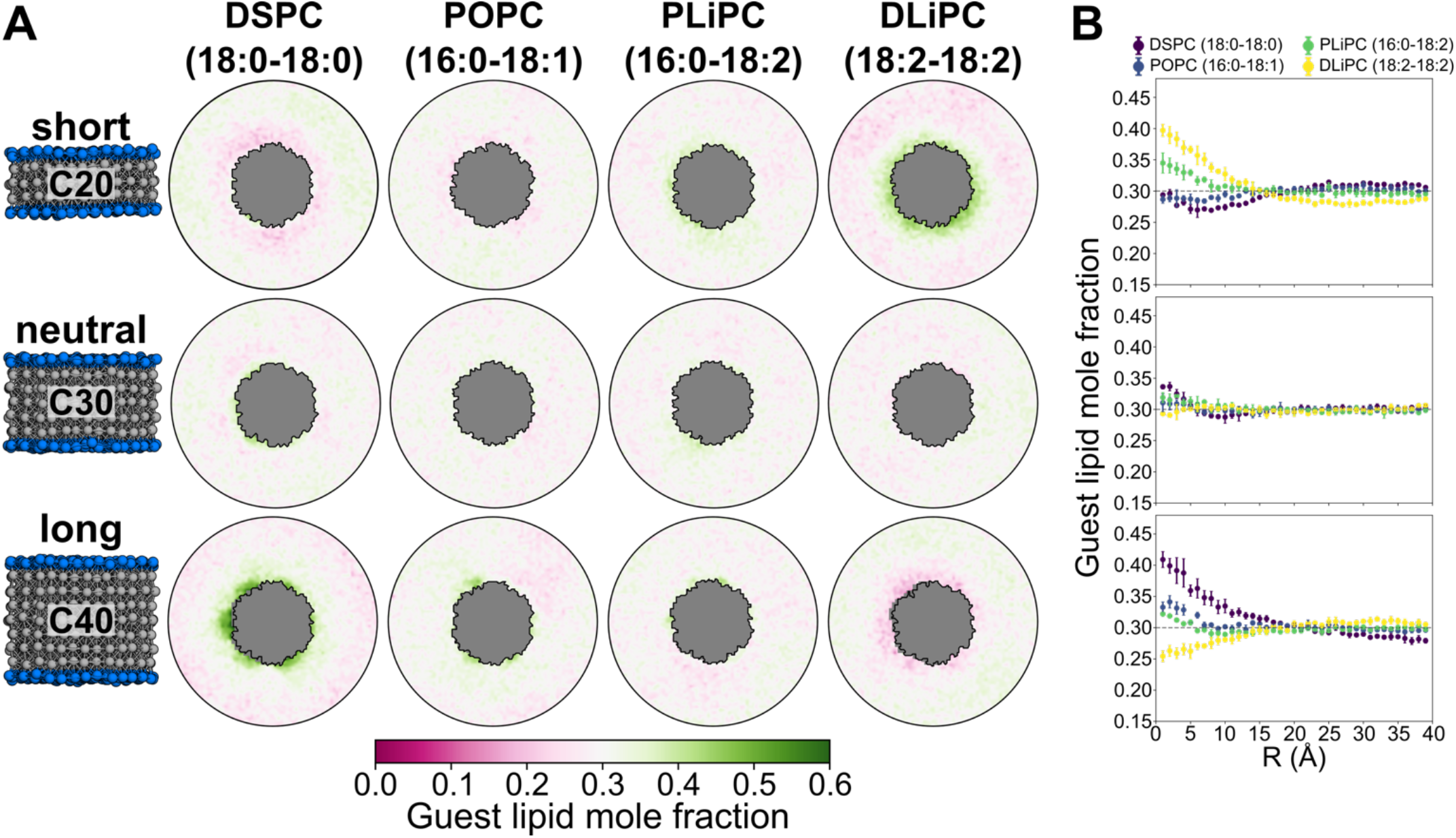
Pseudoproteins change guest lipid distribution based on tail unsaturation. **(a)** Ensemble averaged lipid mole fractions were calculated from aligned and centered CGMD trajectories (1-10 µs) with 30 mol% guest lipid and 70 mol% DOPC (host lipid). Gray represents the pseudoprotein. **(b)** Quantification of data in panel A showing the guest lipid mole fraction as a function of the distance from the protein surface R. Data points are radial averages and standard deviations from block averaging (1-4 µs, 4-7 µs, 7-10 µs).

The enriched mole fractions of DLiPC (18:2-18:2) around C20 and DSPC (18:0-18:0) around C40 were both ~40 mol% at the pseudoprotein surface (*R* = 1 Å), with a concomitant decrease to ~27 mol% far away from the protein (*R* = 40 Å). Using the Boltzmann distribution 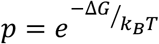 we can translate this equilibrium enrichment to an energy using 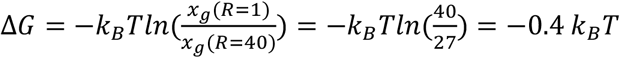. The fact that this result is on the order of a thermal fluctuation underlines the transient “solvent-like” nature of lipids interacting with (protein-induced) membrane thickness deformations (1, 8, 50–53).

### An elastic membrane model reproduces lipid sorting trends from CGMD simulation

Next, we asked if we could reproduce the lipid sorting trends observed in CGMD by minimizing an elastic energy function inspired by the mattress model as a function of the lateral lipid distribution. We parametrized our model with physical membrane properties (hydrophobic thickness *L*_0_, area compressibility *K*_*A*_, bending modulus *K*_*c*_, and area per lipid *A*_lip_) from experimental and all-atom simulation data that were reported in the literature (Table 1). As additional input, we used the leaflet shapes *u*^±^ from our CGMD simulations of pseudoprotein insertions in pure DOPC (Fig. 1b). We initialized (*i* = 0) our model with the bulk lipid fractions (*x*_g,*i*=0_ = *C*_g_ = 0.3; *x*_*h,i*=0_ = 1 − *x*_g,*i*=0_ = 0.7), and a protein radius *R*_*P*_ = 27 Å to match the Van der Waals radii of the pseudoproteins used in CGMD (Fig. 2). Then, for proteins C20, C30, and C40 in each of the four CGMD simulated lipid mixtures, we minimized the free energy using a Markov Chain Monte Carlo (MCMC) scheme that makes random changes in the local guest lipid mole fraction and accepts moves following a Metropolis criterion (see Methods section for details). Ten independent MCMC runs each lasting 5,000 iterations were performed on each lipid system. All MCMC simulations converged within ~1,000 iterations (Fig. 4, Fig. S4, movies in Supporting Material) to typical Δ*G* values of ~ −1 k_B_T, similar to the energies we estimated from the enrichments in CGMD. Again, more deformable unsaturated DLiPC was enriched and the stiffer saturated DSPC was depleted in the thinned vicinity of short-spanning C20 (Fig. 4a,b). Conversely, the thicker membrane around long TM C40 attracted saturated DSPC and repelled unsaturated DLiPC (Fig. 4c,d). Overlaying the converged distributions from our elastic model (see Fig. S5 for the full energy decomposition) with the CGMD data, we see that they are in reasonable agreement with each other (Fig. 5).

**Figure 4.**
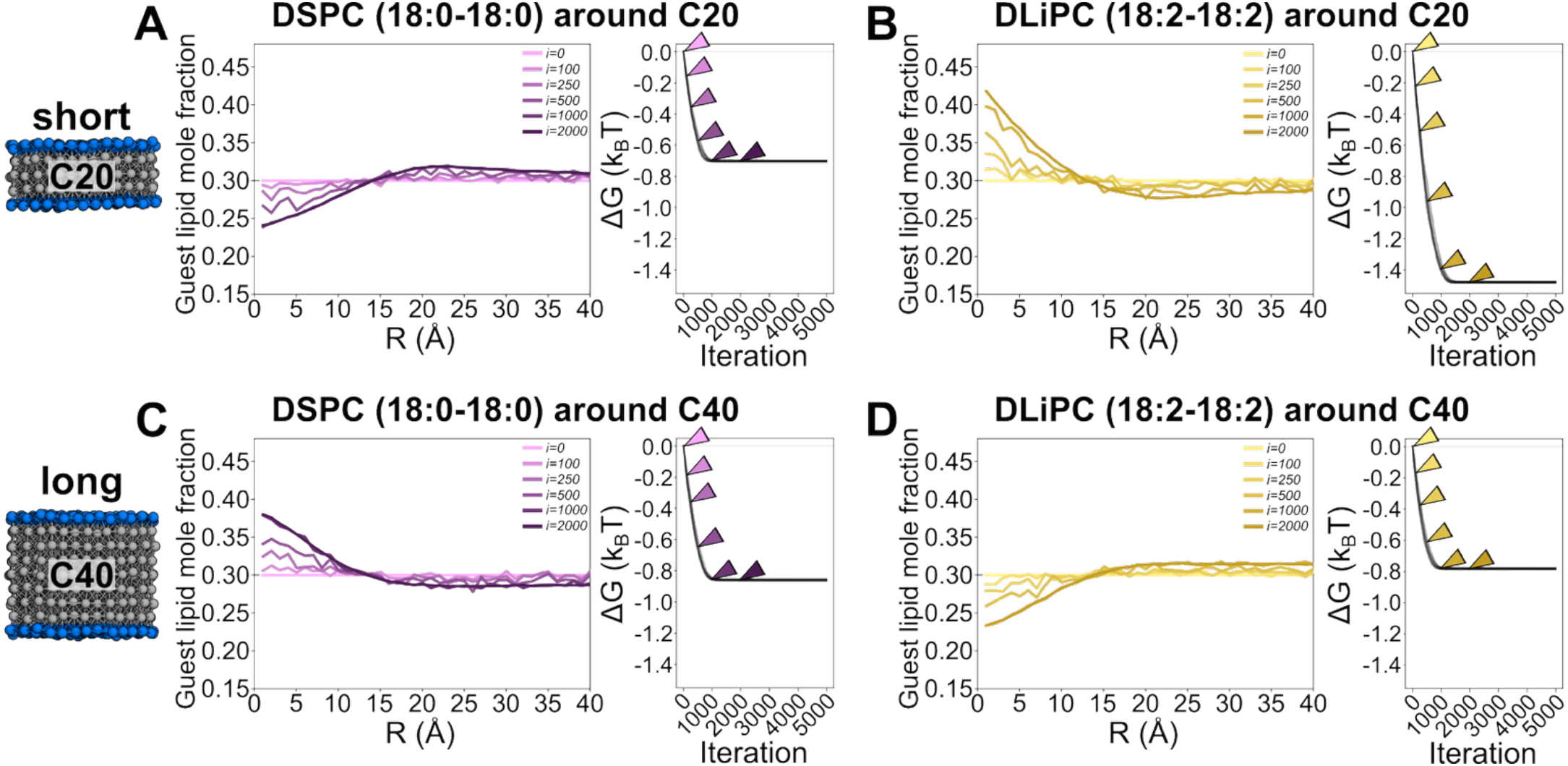
MCMC optimization for DSPC and DLiPC sorting in a continuum elastic membrane model of bilayer compression (C20) and extension (C40). MCMC optimization for DSPC around C20 **(a)**, DLiPC around C20 **(b)**, DSPC around C40 **(c)**, and DLiPC around C40 **(d)**. Left panels: evolution of guest lipid mole fraction as a function of distance from the protein surface for one of the 10 runs. Videos of these processes are provided as Supporting Material. Right panels: Overlay of ΔG minimization curves for ten independent MCMC optimization runs. Colored arrows correspond to individual traces at specific iterations (i) in the respective left-hand panel. DOPC was used as a host lipid (x_h,i=0_ = 0.7) for all calculations. Optimization curves for all guest lipid-pseudoprotein combinations are shown in Fig. S4.

**Figure 5.**
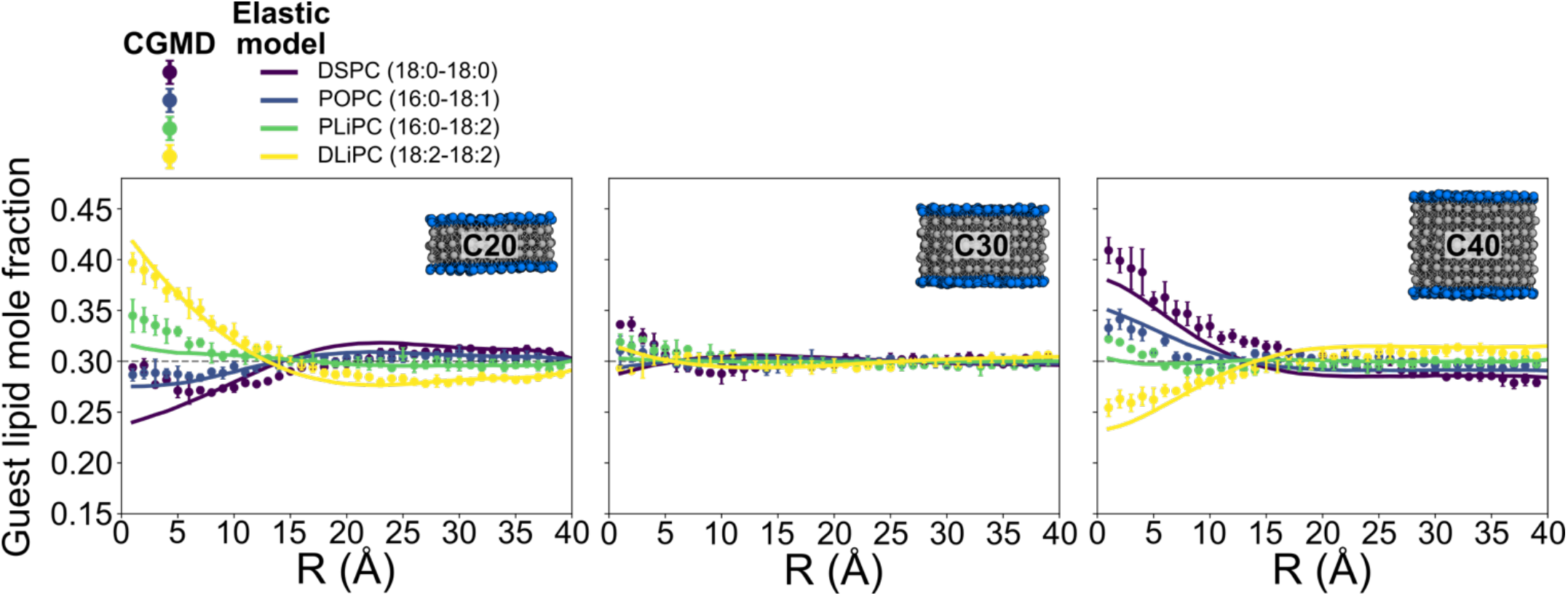
CGMD and the elastic membrane model predict similar lipid sorting trends. CGMD data (circles) are radial averages and standard deviations from block averaging (1-4 µs, 4-7 µs, 7-10 µs, same data as Fig. 3b). Data from our elastic membrane model (lines) are averages and standard deviations (error bars are too small to see) from 10 independent runs with 5,000 iterations each.

We found that the compression energy Δ*H*_comp_ is the dominating term in our energy model driving the enrichment/depletion of the different guest lipid types at the expense of decreasing the mixing entropy (Fig. S5). In contrast, contributions from the bending energy Δ*H*_bend_ to lipid sorting were negligible. There are two explanations for this. First, if we take the C20 protein system as an example, the total compression energy is about four times larger than the bending energy at 40-60 k_B_T *versus* 10-15 k_B_T, respectively (Table S5). This means that there is more compression energy to gain or lose by changing the local lipid composition than there is bending energy. Second, differences between the physical characteristics of the guest lipid and the DOPC host lipid impact the compression energy in two ways: firstly, through modification of the compressibility modulus *K*_*A*_, and secondly by changing the equilibrium lipid *L*_0_ length, which contributes quadratically to the energy (see Eq. 2). In contrast, the only lipid-type dependent term in the bending energy is the modulus *K*_*c*_, the values of which are too similar to contribute to lipid sorting in these calculations.

Because our elastic model is dominated by the compression parameters *L*_0_ and *K*_*A*_, we wondered how sensitive its predictions would be to changes in those values. To probe this, we replaced the physical lipid parameters *L*_0_, *K*_*A*_, *K*_*c*_, and *A*_lip_ from experiments by values obtained from our own CGMD simulations of protein-free single component bilayers (Table S4). CGMD-derived hydrophobic thickness values spanned a larger range compared to the literature values in Table 1: the simulated DSPC bilayer was 1 Å thicker, whereas the simulated DLiPC bilayer was 1.6 Å thinner. Moreover, literature values for the compressibility modulus *K*_*A*_ are very similar for the lipid types considered in this study and all lie between 0.50 to 0.58 k_B_T Å^−2^, whereas values from CGMD box fluctuations were considerably smaller and gradually decreased with increasing tail unsaturation (from 0.41 k_B_T Å^−2^ for DSPC to 0.21 k_B_T Å^−2^ for DLiPC). Interestingly, despite the differences in the length and compressibility between the experimental values and those determined from simulation, the elastic model produced similar results with both sets of parameters (Fig. S6a-d). The similarity in the model predictions occurs because the exaggerated thickness differences obtained from the CGMD parameterization are counter-acted by the lower compression moduli, effectively resulting in similar energies and thus similar lipid sorting. This finding is substantiated by the notion that mixing the literature and simulation parameters (literature *L*_0_ with simulated *K*_*A*_, or *vice versa*) produce strongly divergent lipid distributions that are in poor agreement with our CGMD simulation data (Fig. S6e-h). Taken together, these findings suggest that although the isolated membrane parameters in the Martini 3.0.0 lipid force-field differ from the experimental literature, their combination still produces realistic membrane elastic behavior.

## DISCUSSION

In this paper, we explored how mattress model-like membrane deformations – specifically stretching and thinning the bilayer – impact lipid sorting as a function of tail unsaturation. We performed CGMD simulations of idealized pseudoproteins in two-component membranes where we could control the characteristics of the protein-membrane contact interface across three scenarios adjusting only the protein length. In doing so, we removed as many confounding variables as possible and controlled for protein surface hydrophilicity (C3 Martini bead type) and protein shape (cylindrical, all with the same radii and surface roughness). For the lipids, we kept a constant tail length (4 Martini beads) and headgroup chemistry (PC) and only changed the number of unsaturations in the tails. As a result, we show that *short* TM pseudoproteins attract unsaturated lipids (~33% enrichment) and repel saturated lipids (~15% depletion). Conversely, *long* TM pseudoproteins attract saturated lipids and repel unsaturated species by similar percentages.

We then attempted to match our MD results with an elastic membrane model that balances compression and bending energies with lipid mixing entropy. We parametrized this model with experimental and all-atom simulation data from the literature and applied an MCMC optimization scheme to find the lipid distribution for which the free energy in the two-component membrane is minimal. For most conditions, the model does an excellent job matching the CGMD simulations. This finding indicates that the subtle change in membrane thickness and compressibility when introducing two double bonds to a lipid tail (18:0 to 18:2) coupled with the degree of membrane deformation induced by these proteins is sufficient to drastically change the distribution of lipids around the protein.

Both our CMGD simulations and elastic model produce lipid sorting free energies (Δ*G*) of around 0.5-1 k_B_T, indicating that such states are transient and should be thought of as solvent-like interactions rather than ligand-binding events (1). Although the energies are small, we would like to point out that corresponding enrichment factors in the 1.5-2.5 range are typically considered large enough to surpass the fold-change significance threshold in protein-lipid interactome studies (54). Moreover, it is important to keep in mind that our lipid sorting simulations were done against a background of DOPC, which itself has two double bonds (18:1-18:1). We argue that increasing the difference between the degrees of saturation of the host and guest lipids should exaggerate sorting trends. Moreover, we hypothesize that the energies and resulting lipid enrichment/depletion may be larger in realistic complex membranes that comprise a large variation of tail lengths and degrees of unsaturation, as well as a high concentration of membrane-stiffening cholesterol (18, 55).

### Protein tilt

To be consistent with the original mattress model, we restrained protein tilt in our simulations. However, it is well established in both simulation and NMR studies (56–60) that some proteins – especially long single-pass TM domains – tilt in the membrane to minimize hydrophobic mismatch. To determine how sorting changes with protein tilt, we repeated our simulations without applying a tilt restraint and found that the tilt angle distributions of C20, C30, and C40 in pure DOPC are nearly identical (Fig. S7), with mean angles at a mere ~5º with respect to the z-axis. This modest protein-independent tilt angle likely results from the diameter (54 Å) of our cylinders being larger than the membrane thickness (30 Å), which disfavors tilting (60). We did notice, however, that removing the tilt restraint generally decreased the lipid order near the pseudoproteins (Fig. S8). Interestingly, this concurrently inverted the sorting trends around the long C40 TM protein, such that unsaturated DLiPC is now enriched by ~10% near the protein, while saturated DSPC is depleted by ~10% (Fig. S9). We argue that is due to the unsaturated DLiPC lipids preferring a disordered membrane environment, whereas the saturated lipid tails of DSPC do not. This inversion in sorting trends demonstrates the utility of restraining protein tilt; doing so allowed us to isolate lipid sorting from the confounding lipid tail disorder that protein tilt introduces. The notion that poly-unsaturated lipids are enriched around all pseudoproteins when we allow for protein tilt suggests that there may be a general bias toward accumulating unsaturated species around protein inclusions. This is in line with previous simulation work that showed an accumulation of poly-unsaturated tails near 9 out of 10 assayed proteins, within a complex mixture of 63 lipid types (11). We speculate that this idea may also help explain why many functional lipid molecules (e.g., phosphatidylinositol-4,5-bisphosphate (PIP_2_) (61), arachidonic acid (AA, 20:4), and docosahexaenoic acid (DHA, 22:6)(62)) typically feature acyl tails with multiple double bonds.

### Protein surface roughness

When comparing the CGMD data to our elastic model, there is an interesting mismatch between the two methods for the DSPC-containing mixtures. In CGMD, we observed an uptick in DSPC concentration near the C20, C30, and C40 protein surfaces that was not captured by the elastic model (Fig. 5). This discrepancy likely stems from favorable packing interactions between the saturated DSPC tails and the protein surface that were not accounted for in the elastic model. To explore this further, we performed CGMD simulations of tilt-restrained C40 variants for which we changed the roughness of the pseudoprotein surface by lowering the c_offset_ parameter (0.0 Å; i.e., a smooth, completely flat surface), or doubling the c_offset_ parameter (0.2 Å; i.e., a rougher protein surface); see Fig. S10a. We observed increased lipid order (Fig. S10b) and enrichment of the fully saturated DSPC around the smoother C40 (Fig. S10c) due to a more favorable packing of the straight, ordered, DSPC tails against the flat hydrophobic protein surface. Intriguingly, increasing the roughness of the C40 surface completely abolished the DSPC attraction (Fig. S10c). These results show that lipid tail unsaturation not only determines protein-induced sorting behavior based on the compression/extension of lipid tails, but also through direct protein-lipid interactions beyond typical ligand-like binding modes that often depend on the headgroup chemistry. Indeed, previous studies have found that TM protein inclusions locally reduce lipid order (63) and dynamics, with rougher protein surfaces slowing down the acyl chain mobility more (64).

### What does this mean for real proteins?

Finally, we should discuss how the findings from our highly idealized physical model systems may help us understand real membrane proteins. Sharpe *et al*. showed that the length of TM domains of most eukaryotic proteins has evolved to approximately match the thickness of their respective membrane environment (2). However, their work also describes a large variance in TM length ranging from ~18 to ~32 residues (spanning ~27-48 Å) in the plasma membrane of vertebrates. This implies that there is a considerable fraction of membrane proteins that exhibit hydrophobic mismatch (i.e., they are either too short or too long). Our study shows that these mismatches can change the direct lipid environment, or “proteolipid fingerprint” (11). Despite their transient nature (Δ*G*’s on the order of ~1 k_B_T), we argue that these lipid sorting phenomena can crucially impact protein function and may act synergistically with subsequent lipid binding to well-defined binding sites (e.g., the way PIP_2_ lipids (65) and fatty acids (66) modulate ion channel activity) or by changing the opening/closing threshold of membrane tension-controlled proteins (e.g., how tail length and unsaturation modulate the bacterial mechanosensitive ion channels MscL and MscS (67, 68)). We envision that simplified model systems like the pseudoproteins presented in this work will be valuable to gain fundamental understanding of how membrane proteins affect the local composition and properties of their surrounding lipid membranes.

## DATA AND CODE AVAILABILTY

MD simulation data and MCMC python code are available on Zenodo (DOI: 10.5281/zenodo.20447816).

## ACKNOWLEDGMENTS

This work was supported by grants from the National Science Foundation (MCB 2217662) and National Institutes of Health (R01 GM137109). NvH was supported by a Postdoctoral Fellowship provided by the UCSF CVRI. Lisa Zheng and Jonathan Borowsky are thanked for carefully reading the manuscript. Dr. James Lincoff, Frank Marcoline, and Dr. Charles Wolgemuth provided valuable feedback during the early stages of the project.

## AUTHOR CONTRIBUTIONS

NvH and MG designed the research and developed the theory. NvH conducted and analyzed the simulations. MG obtained the funding. NvH wrote the initial draft of the manuscript and both authors worked on revisions.

## SUPPORTING MATERIAL

Fig. S1 – Pseudoprotein setup

Fig. S2 – Quantifying lipid sorting from CGMD simulations

Fig. S3 – Membrane thickness profiles for two-component lipid mixtures compared to pure DOPC

Fig. S4 – MCMC convergence of the total system’s Δ*G*

Fig. S5 – Energy decomposition

Fig. S6 – Comparison of elastic model results with different input parameters

Fig. S7 – Tilt angle distributions for simulations of C20, C30, and C40 in pure DOPC

Fig. S8 – Lipid order parameter maps around C20, C30, and C40 with and without tilt angle restraints

Fig. S9 – Comparison between lipid sorting in CGMD without tilt angle restraints and elastic model

Fig. S10 – Pseudoprotein surface roughness affects lipid order and sorting of saturated DSPC lipids

Table S1 – Overview of CGMD simulations

Table S2 – Polynomial fitting parameters for membrane shapes

Table S3 – Numerical example of an MCMC update step

Table S4 – Physical lipid parameters derived from pure bilayer CGMD simulations

Table S5 – Total energies before and after MCMC optimization

