## supporting_material for "Hydrophobic mismatch induces lipid sorting based on tail unsaturation"

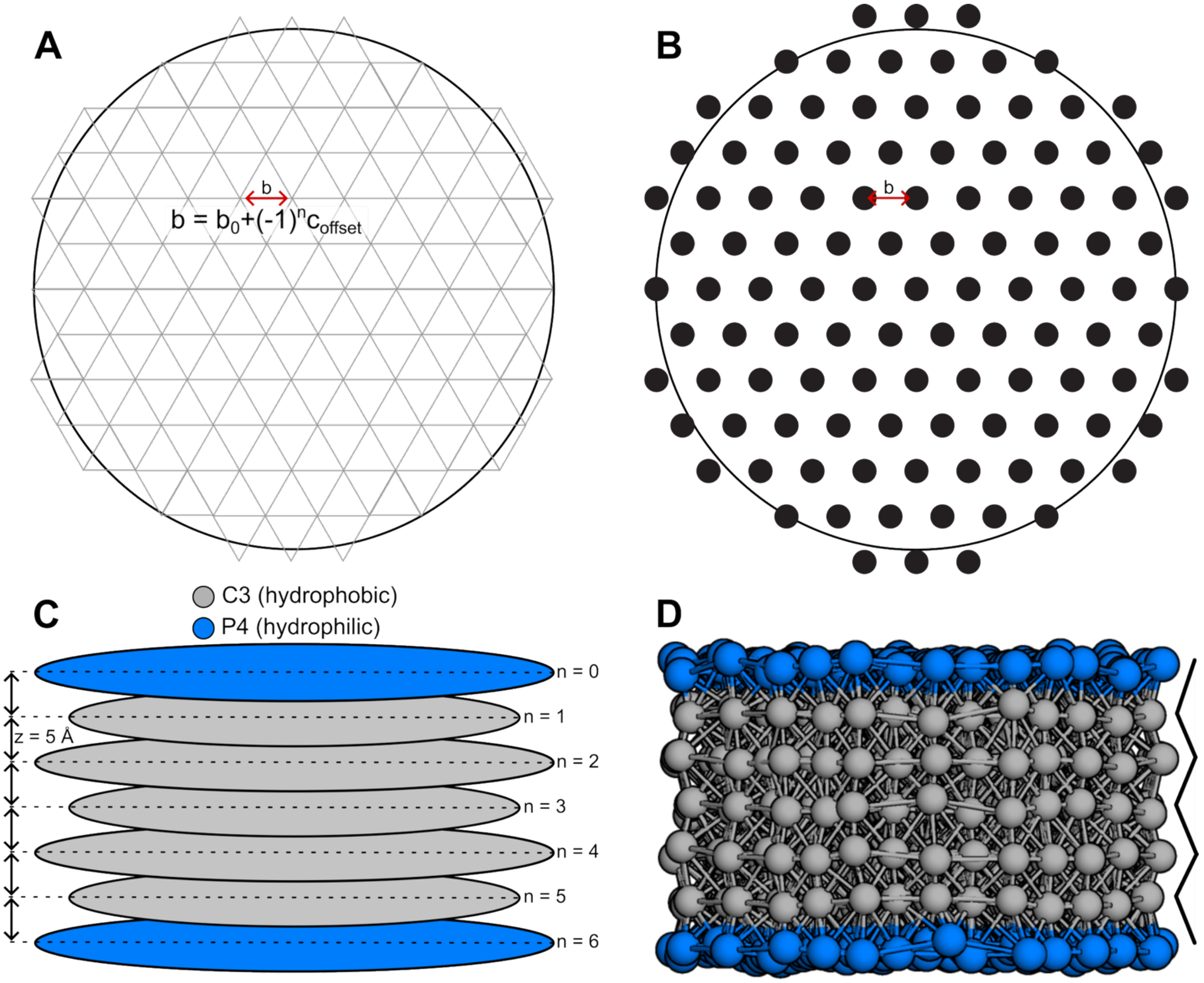

***Figure S1. Pseudoprotein setup. (a)*** *Triangular grid fitted to a circle with a 25 Å radius.* ***(b)*** *Bead placement on each node.* ***(c)*** *Stacked circular layers form a cylinder.* ***(d)*** *3D render of the final fully interconnected C30 pseudoprotein after CGMD equilibration. The zig-zag line highlights the surface roughness, controlled by* $c_{offset}$ *= 0.1 Å.*

***Table S1: Overview of CGMD simulations.*** *Protein-free simulations (top five rows) were conducted to obtain physical parameters $h_0$, $L_0$, $K_A^{app}$, and $A_{lip}$ (Table S4).*

| **Pseudoprotein** | **$c_{offset}$ (Å)** | **Tilt angle restraint** | **Host lipid** | **Guest lipid** | **Initial box size (Å)** | **Simulation time** |
|---|---|---|---|---|---|---|
| - | - | - | DSPC (100%) | - | 250x250x120 | 2 µs |
| - | - | - | POPC (100%) | - | 250x250x120 | 2 µs |
| - | - | - | DOPC (100%) | - | 250x250x120 | 2 µs |
| - | - | - | PLiPC (100%) | - | 250x250x120 | 2 µs |
| - | - | - | DLiPC (100%) | - | 250x250x120 | 2 µs |
| C20 | 0.1 | Yes | DOPC (100%) | - | 140x140x100 | 10 µs |
| C30 | 0.1 | Yes | DOPC (100%) | - | 140x140x100 | 10 µs |
| C40 | 0.1 | Yes | DOPC (100%) | - | 140x140x100 | 10 µs |
| C20 | 0.1 | Yes | DOPC (70%) | DSPC (30%) | 140x140x100 | 10 µs |
| C20 | 0.1 | Yes | DOPC (70%) | POPC (30%) | 140x140x100 | 10 µs |
| C20 | 0.1 | Yes | DOPC (70%) | PLiPC (30%) | 140x140x100 | 10 µs |
| C20 | 0.1 | Yes | DOPC (70%) | DLiPC (30%) | 140x140x100 | 10 µs |
| C30 | 0.1 | Yes | DOPC (70%) | DSPC (30%) | 140x140x100 | 10 µs |
| C30 | 0.1 | Yes | DOPC (70%) | POPC (30%) | 140x140x100 | 10 µs |
| C30 | 0.1 | Yes | DOPC (70%) | PLiPC (30%) | 140x140x100 | 10 µs |
| C30 | 0.1 | Yes | DOPC (70%) | DLiPC (30%) | 140x140x100 | 10 µs |
| C40 | 0.1 | Yes | DOPC (70%) | DSPC (30%) | 140x140x100 | 10 µs |
| C40 | 0.1 | Yes | DOPC (70%) | POPC (30%) | 140x140x100 | 10 µs |
| C40 | 0.1 | Yes | DOPC (70%) | PLiPC (30%) | 140x140x100 | 10 µs |
| C40 | 0.1 | Yes | DOPC (70%) | DLiPC (30%) | 140x140x100 | 10 µs |
| C20 | 0.1 | No | DOPC (100%) | - | 140x140x100 | 10 µs |
| C30 | 0.1 | No | DOPC (100%) | - | 140x140x100 | 10 µs |
| C40 | 0.1 | No | DOPC (100%) | - | 140x140x100 | 10 µs |
| C20 | 0.1 | No | DOPC (70%) | DSPC (30%) | 140x140x100 | 10 µs |
| C20 | 0.1 | No | DOPC (70%) | POPC (30%) | 140x140x100 | 10 µs |
| C20 | 0.1 | No | DOPC (70%) | PLiPC (30%) | 140x140x100 | 10 µs |
| C20 | 0.1 | No | DOPC (70%) | DLiPC (30%) | 140x140x100 | 10 µs |
| C30 | 0.1 | No | DOPC (70%) | DSPC (30%) | 140x140x100 | 10 µs |
| C30 | 0.1 | No | DOPC (70%) | POPC (30%) | 140x140x100 | 10 µs |
| C30 | 0.1 | No | DOPC (70%) | PLiPC (30%) | 140x140x100 | 10 µs |
| C30 | 0.1 | No | DOPC (70%) | DLiPC (30%) | 140x140x100 | 10 µs |
| C40 | 0.1 | No | DOPC (70%) | DSPC (30%) | 140x140x100 | 10 µs |
| C40 | 0.1 | No | DOPC (70%) | POPC (30%) | 140x140x100 | 10 µs |
| C40 | 0.1 | No | DOPC (70%) | PLiPC (30%) | 140x140x100 | 10 µs |
| C40 | 0.1 | No | DOPC (70%) | DLiPC (30%) | 140x140x100 | 10 µs |
| C40 | 0.0 | Yes | DOPC (70%) | DSPC (30%) | 140x140x100 | 10 µs |
| C40 | 0.2 | Yes | DOPC (70%) | DSPC (30%) | 140x140x100 | 10 µs |

***Table S2: Polynomial fitting parameters for membrane shapes.*** *We fit a 6th degree polynomial to the averaged GL1 and GL2 coordinates between* $R = 1$ Å *and* $R = 40$ Å *from pure DOPC CGMD simulations around each pseudoprotein to get* $u(r)$ *in the form:* $u(r) = c_1 r^5 + c_2 r^4 + c_3 r^3 + c_4 r^2 + c_5 r + c_6$, *with* $r = R + R_P$ *and* $u(r) = 0$ *as the bilayer midplane.*

| Pseudoprotein | Leaflet | $c_1$ | $c_2$ | $c_3$ | $c_4$ | $c_5$ | $c_6$ |
|---|---|---|---|---|---|---|---|
| C20 | upper ($u^+$) | $-2.649 \times 10^{-7}$ | $6.347 \times 10^{-5}$ | $-5.801 \times 10^{-3}$ | $2.464 \times 10^{-1}$ | $-4.607$ | $40.47$ |
| C20 | lower ($u^-$) | $2.743 \times 10^{-7}$ | $-6.578 \times 10^{-5}$ | $6.024 \times 10^{-3}$ | $-2.571 \times 10^{-1}$ | $4.857$ | $-42.75$ |
| C30 | upper ($u^+$) | $-1.133 \times 10^{-8}$ | $1.575 \times 10^{-6}$ | $1.453 \times 10^{-5}$ | $-1.154 \times 10^{-2}$ | $5.951 \times 10^{-1}$ | $5.805$ |
| C30 | lower ($u^-$) | $8.321 \times 10^{-9}$ | $-6.595 \times 10^{-7}$ | $-1.168 \times 10^{-4}$ | $1.691 \times 10^{-2}$ | $-7.294 \times 10^{-1}$ | $-4.525$ |
| C40 | upper ($u^+$) | $2.249 \times 10^{-7}$ | $-5.573 \times 10^{-5}$ | $5.347 \times 10^{-3}$ | $-2.447 \times 10^{-1}$ | $5.206$ | $-24.06$ |
| C40 | lower ($u^-$) | $-2.184 \times 10^{-7}$ | $5.446 \times 10^{-5}$ | $-5.259 \times 10^{-3}$ | $2.424 \times 10^{-1}$ | $-5.192$ | $24.25$ |

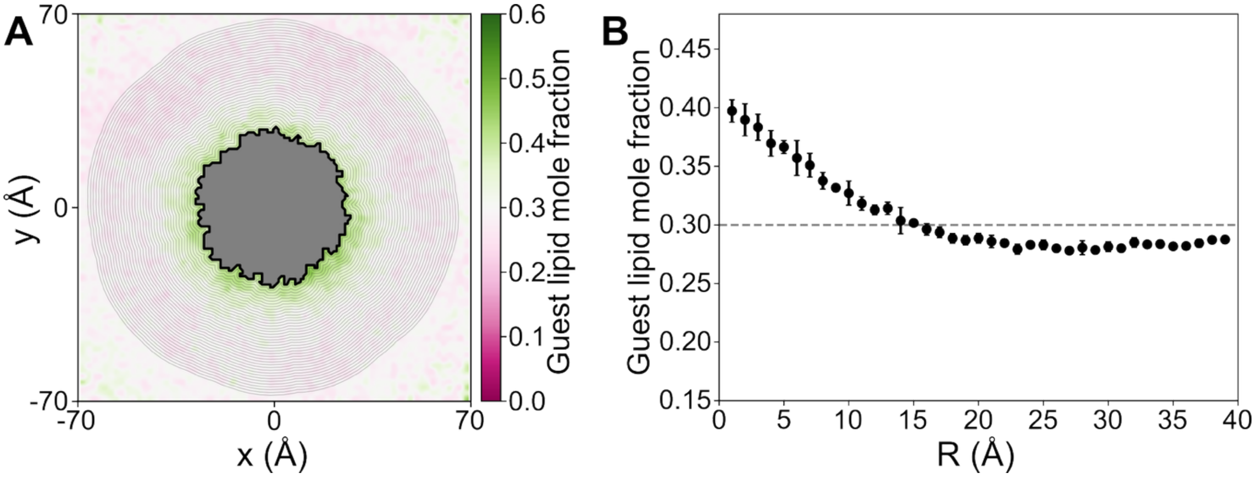

***Figure S2. Quantifying lipid sorting from CGMD simulations. (a)*** *Ensemble-averaged map of the guest lipid mole fraction around C20 in the DLiPC:DOPC (3:7) lipid mixture. Isosurface bins with a 1 Å bin width were drawn around the pseudoprotein shape.* ***(b)*** *Radial averages and standard deviations from block averaging (1-4 μs, 4-7 μs, 7-10 μs). Each data point corresponds to the average mole fraction in a bin between two consecutive isosurface lines in panel a.*

***Table S3: Numerical example of an MCMC update step.*** *A simplified example of the first step in the MCMC scheme for a system with 5 bins. Numbers marked in yellow ($\alpha_{\text{rand}}$) are randomly generated from a uniform distribution between -0.5 and 0.5*. The green number ($\alpha_{\text{final}} = -\sum_{j=1}^{N-1} \alpha_{\text{rand}}$) is chosen to balance the yellow numbers such that $\sum_{j=1}^{N} \alpha = 0$ ensuring that the total number of lipids in the system remains constant. For this example, we used $A_{\text{lip}} = 68.16$ Å$^2$ and $R_P = 27$ Å. The energy numbers are toy examples.*

| ***Description*** | | | | | | $\boldsymbol{N-1}$ | $\boldsymbol{N}$ | |
|---|---|---|---|---|---|---|---|---|
| *Bin index* | $\boldsymbol{j}$ | | **1** | **2** | **3** | **4** | **5** | $\sum_{j=1}^{N}$ |
| *Distance to protein surface (Å)* | $\boldsymbol{R_j}$ | | 1 | 2 | 3 | 4 | 5 | |
| *Guest lipid mole fraction at $i = 0$* | $\boldsymbol{x_{g,i=0,j}}$ | $C_g$ | 0.30 | 0.30 | 0.30 | 0.30 | 0.30 | |
| *Area of radial bin (Å$^2$)* | $\boldsymbol{A_j}$ | $2\pi d(R_j + R_P)$ | 175.9 | 182.2 | 188.5 | 194.8 | 201.1 | |
| *Total lipid number* | $\boldsymbol{M_j}$ | $2A_j/A_{lip}$ | 5.41 | 5.61 | 5.80 | 5.99 | 6.19 | 29.00 |
| *Guest lipid number at $i = 0$* | $\boldsymbol{n_{g,i=0,j}}$ | $x_{g,i=0,j}M_j$ | 1.62 | 1.68 | 1.74 | 1.80 | 1.86 | 8.70 |
| *Random change in guest lipid number between -0.5 and 0.5** | $\boldsymbol{\alpha_j}$ | | 0.50 | -0.24 | -0.49 | -0.11 | 0.34 | 0.00 |
| *Updated guest lipid number at $i = 1$* | $\boldsymbol{n_{g,i=1,j}}$ | $n_{g,i=0,j} + \alpha_j$ | 2.12 | 1.44 | 1.25 | 1.69 | 2.19 | 8.70 |
| *Updated guest lipid mole fraction at $i = 1$* | $\boldsymbol{x_{g,i=1,j}}$ | $n_{g,i=1,j}/M_j$ | 0.39 | 0.26 | 0.22 | 0.28 | 0.35 | |
| ***Toy energy numbers*** | | | | | | | | |
| *Free energy at $i = 0$ ($k_B T$)* | $\boldsymbol{G(R_j, x_{g,i=0,j})}$ | $H_{comp} + H_{bend} - TS_{mix}$ | -0.5 | -0.2 | 0.0 | 0.3 | 0.2 | -0.2 |
| *Free energy difference at $i = 0$ ($k_B T$)* | $\Delta\boldsymbol{G(x_{g,i=0})}$ | $\sum_{j=1}^{N}[G(R_j, x_{g,i=0,j}) - G(R_j, x_{g,i=0,j})]$ | | | | | | 0.0 |
| *Free energy at $i = 1$ ($k_B T$)* | $\boldsymbol{G(R_j, x_{g,i=1,j})}$ | $H_{comp} + H_{bend} - TS_{mix}$ | -1.2 | 0.4 | -0.1 | 0.3 | -0.1 | -0.7 |
| *Free energy difference at $i = 1$ ($k_B T$)* | $\Delta\boldsymbol{G(x_{g,i=1})}$ | $\sum_{j=1}^{N}[G(R_j, x_{g,i=1,j}) - G(R_j, x_{g,i=0,j})]$ | | | | | | -0.5 |
| *Acceptance probability* | $\boldsymbol{P_i}$ | $e^{\frac{\Delta G(x_{g,i=0}) - \Delta G(x_{g,i=1})}{k_B T}}$ | | | | | | 1.65 |
| | | | | | | | | $P_i \geq 1$; the update is accepted |

* We used random numbers between -0.01 and 0.01 in the actual MCMC optimization.

***Table S4: Physical lipid parameters derived from pure bilayer CGMD simulations.*** *All simulations were carried out at 323 K. All values are averages over last 1 μs of the simulation. Hydrophobic thickness* $L_0$ *was measured as the peak-to-peak distance of the glycerol (GL1 and GL2 beads) density plot. The apparent area compressibility modulus* $K_A^{app} = \frac{a_0}{\langle (a-a_0)^2 \rangle} k_B T$ *was obtained from box fluctuations around the mean box area* $a_0$*. We applied the membrane undulation correction proposed by Waheed and Edholm (1) to obtain the "true" compressibility moduli* $\frac{1}{K_A} = \frac{1}{K_A^{app}} - \frac{N_{lip} A_{lip} k_B T}{16.6\pi^3 K_c^2}$*, where* $N_{lip} = 879$ *(the number of lipids in a monolayer), and* $K_c$ *is the bending modulus from experiments (Table 1 in main text).*

| **Lipid** | **Martini abbr.** | $L_0$ **(Å)** | $A_{lip}$ **(Å$^2$)** | $K_A^{app}$ **($k_B$T Å$^{-2}$)** | $K_A$ **($k_B$T Å$^{-2}$)** |
|---|---|---|---|---|---|
| DSPC (18:0-18:0) | DPPC | 32.98 | 61.76 | 0.39 | 0.41 |
| POPC (16:0-18:1) | POPC | 30.14 | 67.65 | 0.35 | 0.38 |
| DOPC (18-1:18:1) | DOPC | 28.99 | 70.20 | 0.32 | 0.36 |
| PLiPC (16:0-18:2) | PIPC | 28.49 | 71.30 | 0.26 | 0.33 |
| DLiPC (18:2-18:2) | DIPC | 25.82 | 77.55 | 0.17 | 0.21 |

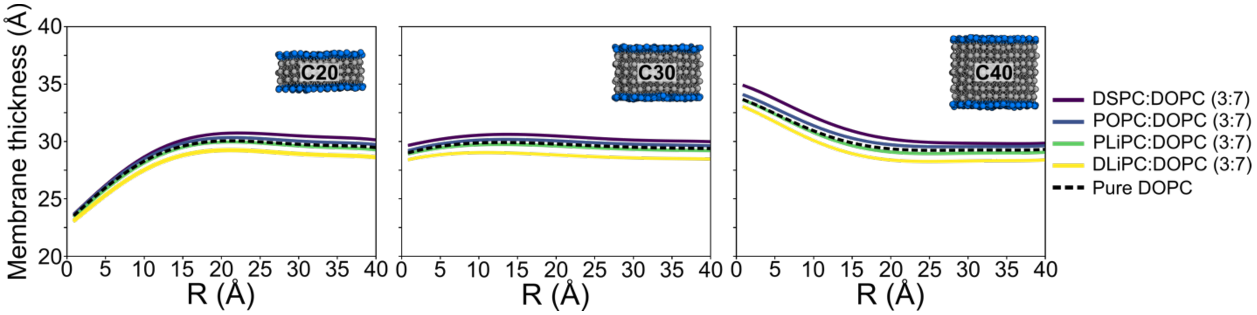

***Figure S3. Membrane thickness profiles for two-component lipid mixtures compared to pure DOPC.***

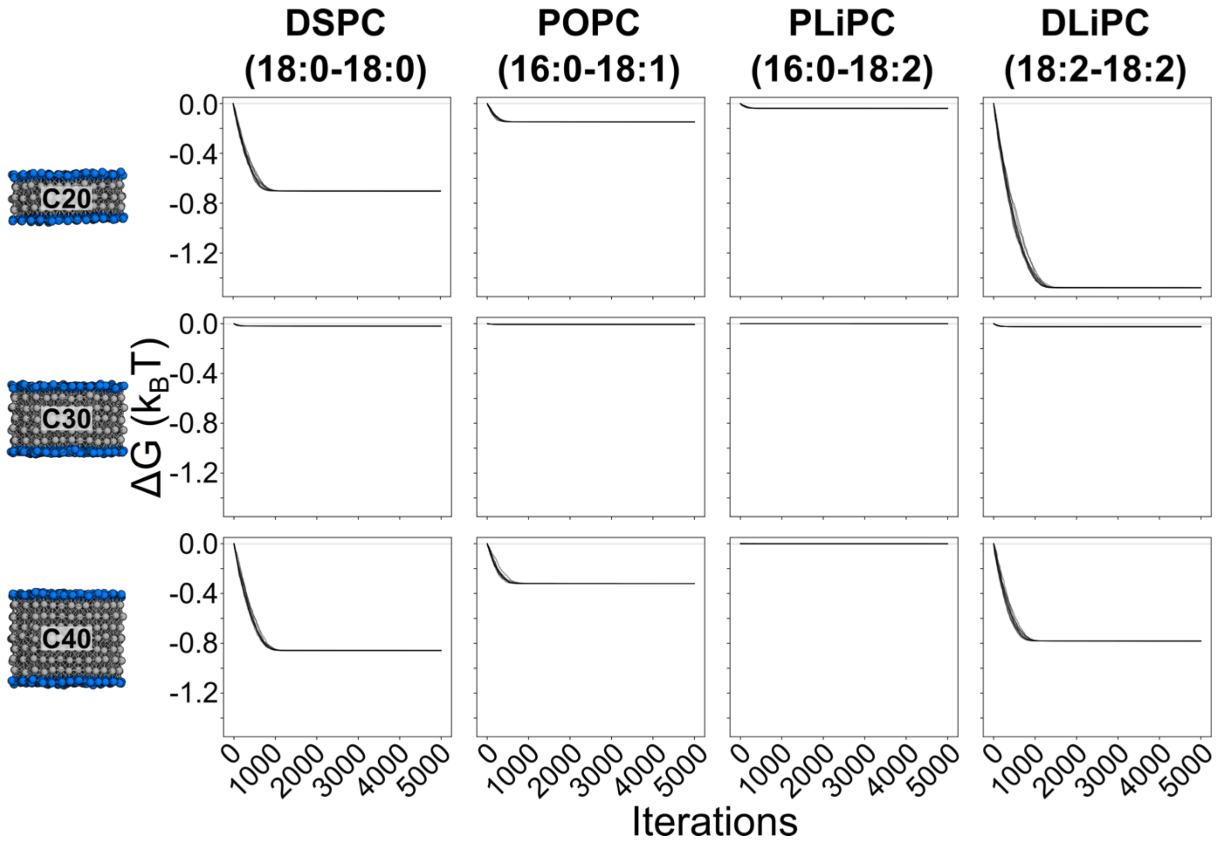

***Figure S4. MCMC convergence of the total system's $\Delta G$ (n=10).***

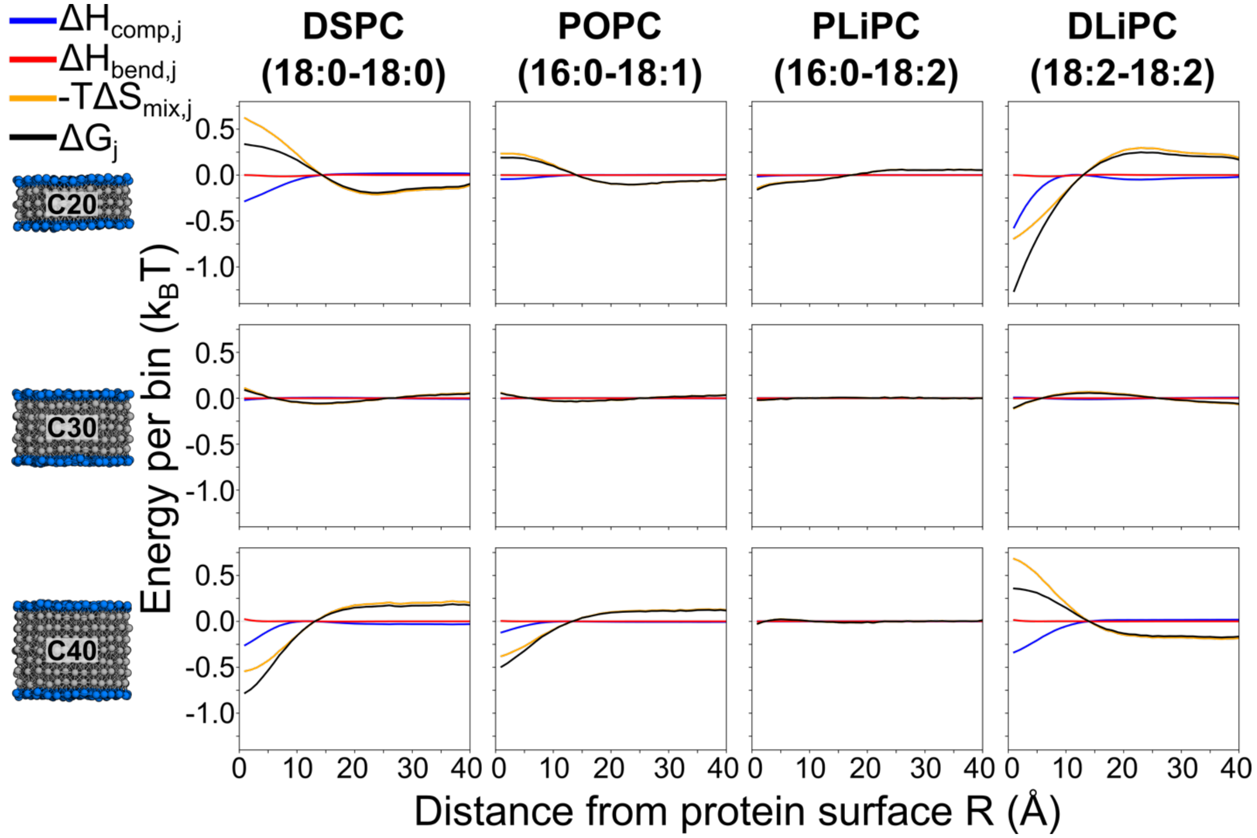

***Figure S5. Energy decomposition.*** *Average energy per bin after 5,000 MCMC iterations (n=10). Error bars are too small to see.*

***Table S5: Total energies before and after MCMC optimization.*** *Compression, bending, and mixing entropy summed over all bins before lipid mixing (3:7 everywhere,* $i = 0$*) and after lipid mixing (MCMC-optimized lipid compositions,* $i = 5000$*), averaged over ten independent MCMC runs. Standard deviations are all smaller than 0.01* $k_BT$*.*

| | | **Compression energy ($k_BT$)** $\sum_{j=1}^{N} H_{\mathrm{comp}}$ | | **Bending energy ($k_BT$)** $\sum_{j=1}^{N} H_{\mathrm{bend}}$ | | **Mixing entropy ($k_BT$)** $T\sum_{j=1}^{N} S_{\mathrm{mix}}$ | |
|---|---|---|---|---|---|---|---|
| **Pseudoprotein** | **Lipid mixture** | Before ($i = 0$) | After ($i = 5000$) | Before ($i = 0$) | After ($i = 5000$) | Before ($i = 0$) | After ($i = 5000$) |
| C20 | DSPC:DOPC (3:7) | 58.7 | 57.4 | 15.5 | 15.4 | 340.9 | 340.3 |
| C20 | POPC:DOPC (3:7) | 43.2 | 42.9 | 14.1 | 14.0 | 340.2 | 340.1 |
| C20 | PLiPC:DOPC (3:7) | 34.0 | 33.9 | 11.7 | 11.7 | 343.9 | 343.9 |
| C20 | DLiPC:DOPC (3:7) | 43.3 | 40.3 | 11.8 | 11.8 | 333.1 | 331.6 |
| C30 | DSPC:DOPC (3:7) | 17.4 | 17.4 | 2.0 | 2.0 | 340.9 | 340.9 |
| C30 | POPC:DOPC (3:7) | 5.6 | 5.6 | 1.8 | 1.8 | 340.2 | 340.2 |
| C30 | PLiPC:DOPC (3:7) | 0.7 | 0.7 | 1.5 | 1.5 | 343.9 | 343.9 |
| C30 | DLiPC:DOPC (3:7) | 18.3 | 18.3 | 1.5 | 1.5 | 333.1 | 333.1 |
| C40 | DSPC:DOPC (3:7) | 31.1 | 29.4 | 5.7 | 5.7 | 340.9 | 340.1 |
| C40 | POPC:DOPC (3:7) | 21.9 | 21.2 | 5.2 | 5.2 | 340.2 | 340.0 |
| C40 | PLiPC:DOPC (3:7) | 21.3 | 21.3 | 4.3 | 4.3 | 343.9 | 343.9 |
| C40 | DLiPC:DOPC (3:7) | 45.1 | 43.6 | 4.3 | 4.4 | 333.1 | 332.4 |

**A**

| | Lit. $L_0$ (Å) | Lit. $K_A$ ($k_BT$ Å$^{-2}$) |
|---|---|---|
| DSPC | 31.98 | 0.54 |
| POPC | 30.89 | 0.52 |
| DOPC | 29.57 | 0.58 |
| PLiPC | 29.40 | 0.54 |
| DLiPC | 27.43 | 0.50 |

**C**

| | Sim. $L_0$ (Å) | Sim. $K_A$ ($k_BT$ Å$^{-2}$) |
|---|---|---|
| DSPC | 32.98 | 0.41 |
| POPC | 30.14 | 0.38 |
| DOPC | 28.99 | 0.36 |
| PLiPC | 28.49 | 0.33 |
| DLiPC | 25.82 | 0.21 |

**E**

| | Lit. $L_0$ (Å) | Sim. $K_A$ ($k_BT$ Å$^{-2}$) |
|---|---|---|
| DSPC | 31.98 | 0.41 |
| POPC | 30.89 | 0.38 |
| DOPC | 29.57 | 0.36 |
| PLiPC | 29.40 | 0.33 |
| DLiPC | 27.43 | 0.21 |

**G**

| | Sim. $L_0$ (Å) | Lit. $K_A$ ($k_BT$ Å$^{-2}$) |
|---|---|---|
| DSPC | 32.98 | 0.54 |
| POPC | 30.14 | 0.52 |
| DOPC | 28.99 | 0.58 |
| PLiPC | 28.49 | 0.54 |
| DLiPC | 25.82 | 0.50 |

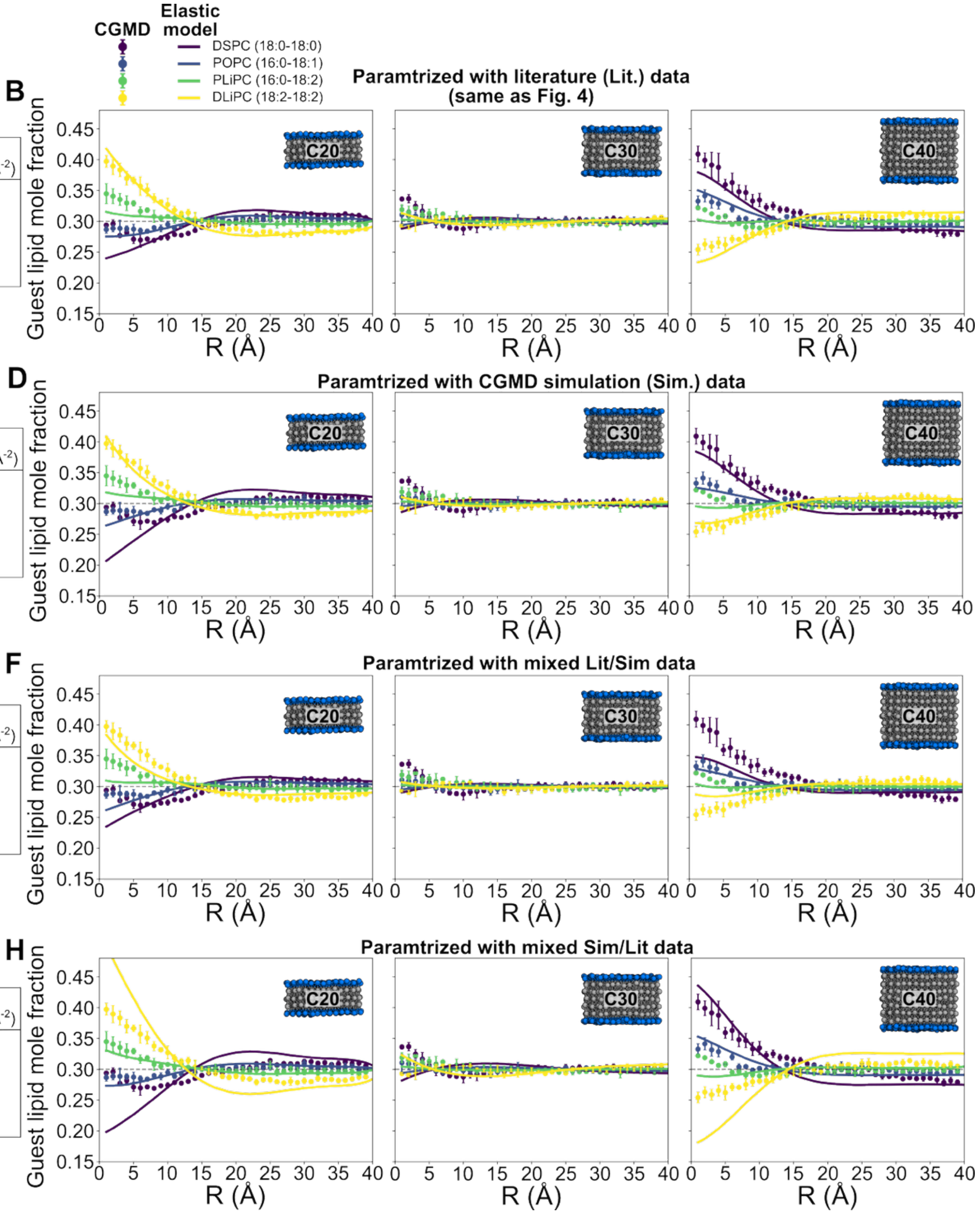

***Figure S6. Comparison of elastic model results with different input parameters. (a, b)*** *Elastic model parameters (a) and results (b), if parametrized with experimental literature (Lit.) data (same as in Table 1). Panel b is the same as Fig. 5 in the main text.* ***(c, d)*** *Elastic model parameters (c) and results (d), if parametrized with CGMD simulation (Sim.) data.* ***(e, f)*** *Elastic model parameters (e) and results (f), if using a mixed parameter set with literature* $L_0$ *and* $K_A$ *from CGMD simulations.* ***(g, h)*** *Elastic model parameters (g) and results (h), if using a mixed parameter set with* $L_0$ *from CGMD simulations and* $K_A$ *from the literature. The CGMD data (solid circles) in panels b, d, f, and h are identical to the data in Fig. 3b in the main text.*

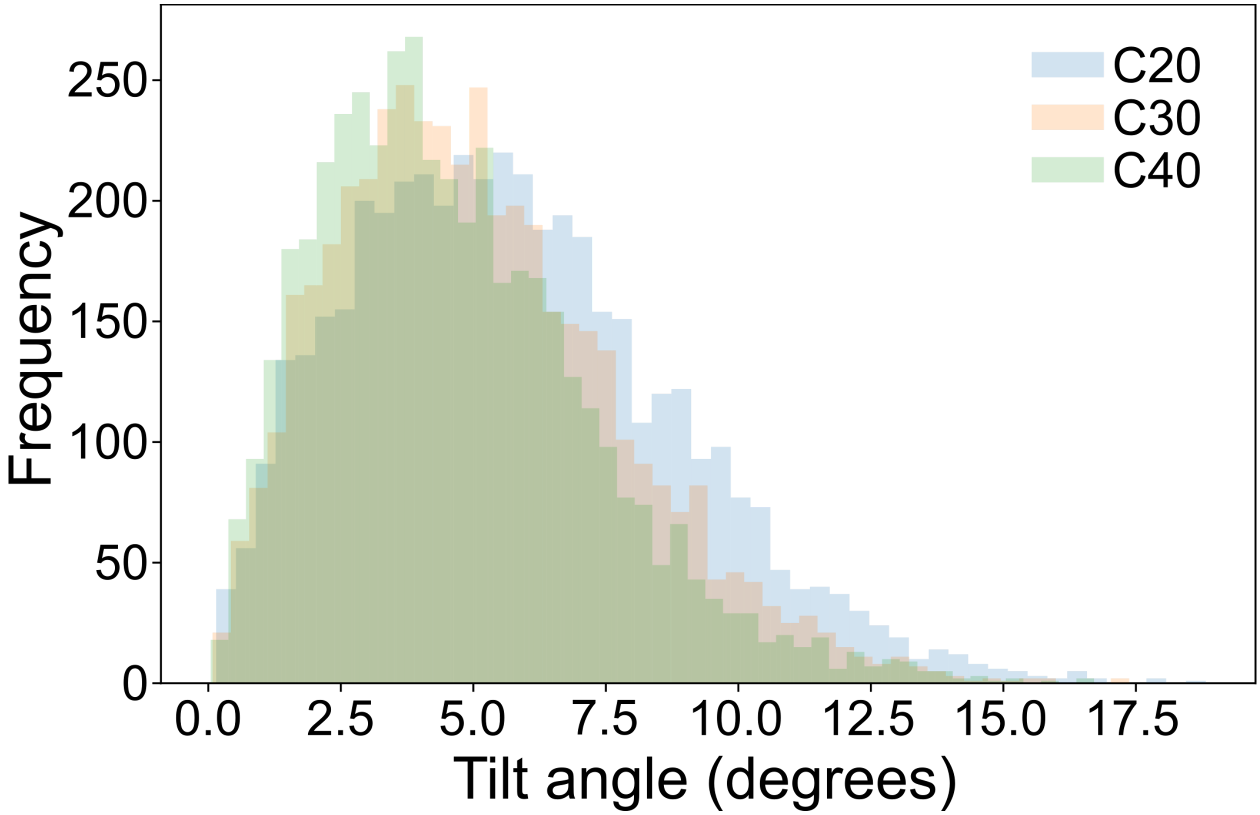

***Figure S7. Tilt angle distributions for simulations of C20, C30, and C40 in pure DOPC.*** *CGMD simulations were carried out without a tilt angle restraint.*

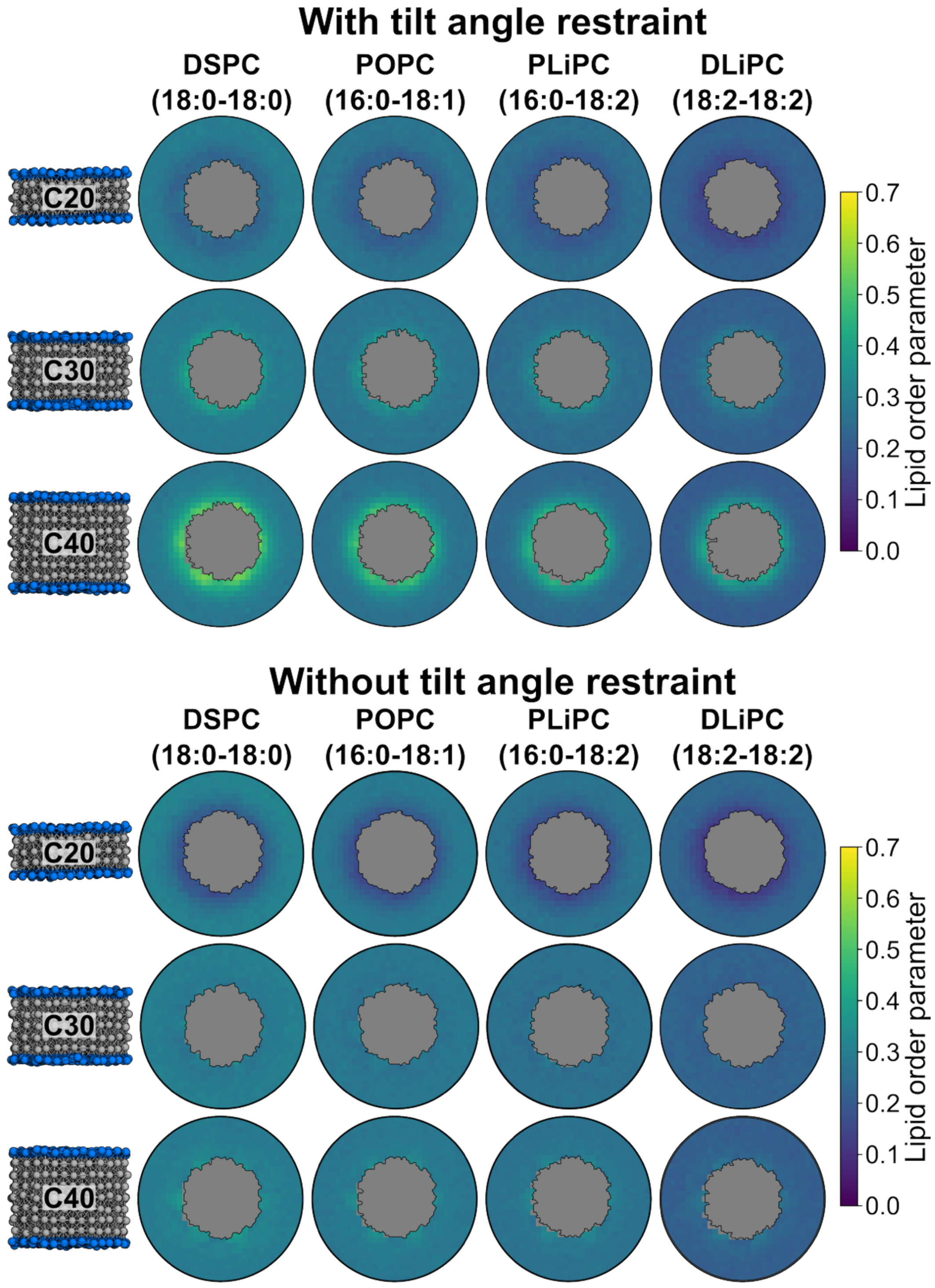

***Figure S8. Lipid order parameter maps around C20, C30, and C40 with (top) and without (bottom) tilt angle restraints.***

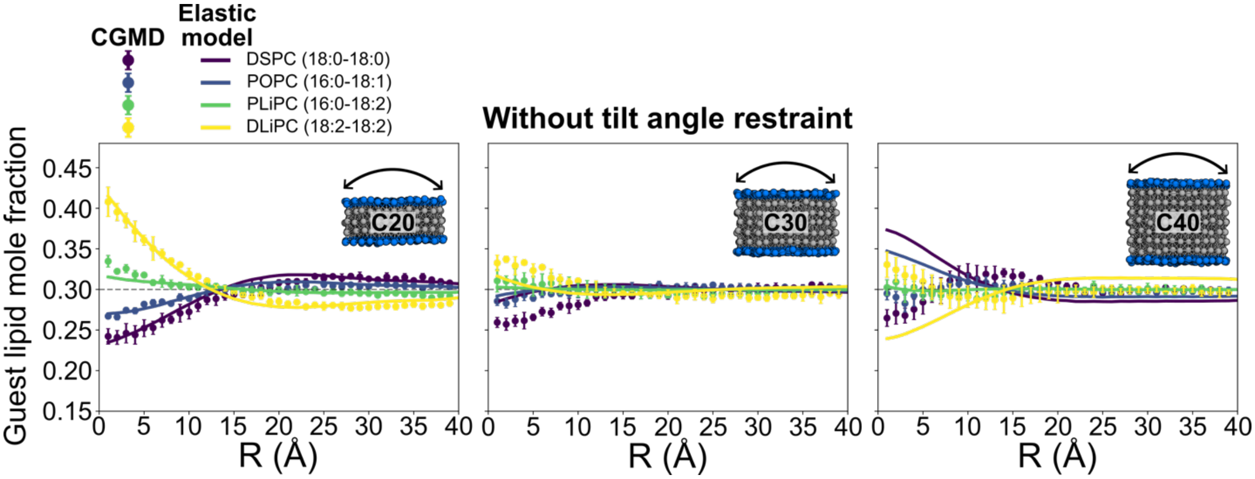

***Figure S9. Comparison between lipid sorting in CGMD without tilt angle restraints and elastic model.*** *CGMD data (circles) are radial averages and standard deviations from block averaging (1-4 μs, 4-7 μs, 7-10 μs). Elastic model data (lines) are averages and standard deviations (error bars are too small to see) from 10 independent runs with 5,000 iterations each.*

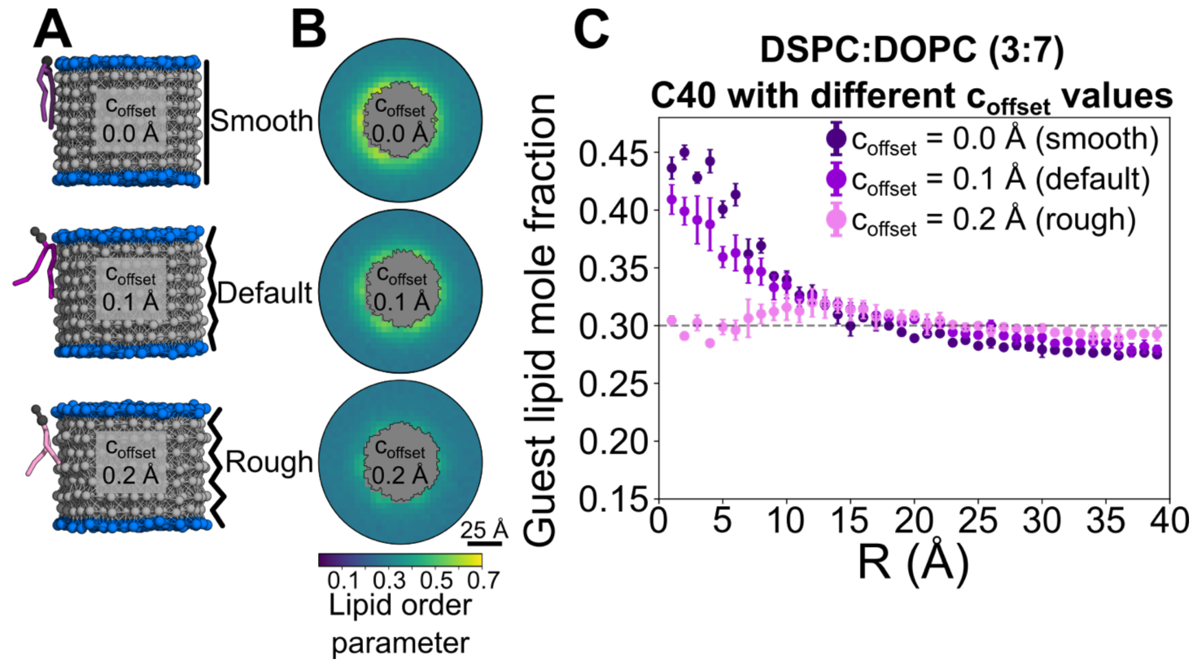

***Figure S10. Pseudoprotein surface roughness affects lipid order and sorting of saturated DSPC lipids. (a)*** *CGMD simulation snapshots of representative DSPC lipids adhered to the surface of C40 pseudoproteins with different* $c_{offset}$ *values to control surface roughness. The default value for all other figures in this paper is* $c_{offset} = 0.1$ *Å.* ***(b)*** *Ensemble-averaged lipid order parameter maps around C40 with different* $c_{offset}$ *values in DSPC:DOPC (3:7) mixtures.* ***(c)*** *Radial averages and standard deviations of guest (DSPC) lipid mole fraction around C40 with different* $c_{offset}$ *values from CGMD simulation.*
